# Encoding neuronal shape in the stochastic dynamics of branching processes

**DOI:** 10.64898/2026.06.05.729577

**Authors:** Marc-Eric Perrin, Maximilien Courgeon, Elise da Silva, Jean-Marc Philippe, Jean-François Rupprecht, Claire Bertet, Thomas Lecuit

## Abstract

Cell shape critically influences function, yet how complex and reproducible morphologies emerge from stochastic cellular dynamics remains unclear. Here, we investigate dendritic morphogenesis of two classes of *Drosophila* mechanosensory neurons with contrasting architectures, combining *in vivo* live imaging, quantitative analysis, cytoskeletal perturbations, and computational modeling. We show that despite sharing similar local stochastic branching rules, the two classes exhibit divergent growth dynamics that cannot be explained by standard, diffusive growth models. This discrepancy arises because Class I neurons display subdiffusive branch dynamics over long timescales, unlike Class IV. Based on these findings, we develop a minimal model with only four parameters that separates short-and long-term branch behaviors, and successfully recapitulates growth dynamics and final morphologies in both classes. Cytoskeletal perturbations reveal a functional separation between actin, which drives short-term exploratory branch fluctuations and arbor expansion, and microtubules, which tune long-term branch diffusivity and determine class-specific morphology. Together, these results establish a parsimonious, generalizable framework linking local cytoskeletal regulation to global neuronal architecture and reveal how stochastic dynamics encode reproducible cell shapes.

## Introduction

Across evolution, organisms have diversified into a remarkable range of forms, from simple body plans to highly elaborate architectures. This diversity extends to the cellular scale, where form and function are tightly coupled. In developing embryos, morphogenesis arises from the interplay between deterministic programs and stochastic fluctuations^1^, as exemplified during branching morphogenesis in organs such as the mammary gland, kidney and lung and the vascular system^2–7^. How these mechanisms collectively generate reproducible, functionally optimized, and often highly complex single cell morphologies remains a fundamental question.

Neurons provide an ideal model to explore this question. They exhibit unparalleled morphological diversity^8^, and their dendritic arbors, typically elaborate and extensively branched, define how signals are received, integrated and processed (Branco and Häusser 2010; London and Häusser 2005). Dendrite morphogenesis is a highly dynamic process involving iterative local branch behaviors (Dailey and Smith 1996; Ferreira Castro et al. 2020; Fujishima, Kawabata Galbraith, and Kengaku 2018; Palavalli et al. 2021; Smith et al. 2010; Wu and Cline 2003) — including elongation, retraction, branching, stabilization and contact-induced self-repulsion —driven by cytoskeletal remodeling^19,20^. Given the importance of dendritic architecture for neuronal function, this process has long been envisioned as predominantly deterministic, governed by intrinsic transcriptional programs and extracellular cues^21,22^.

Recent studies combining live imaging and computational modeling have challenged this long-standing view, by revealing that local branch dynamics play a central, previously underappreciated role in shaping dendritic arbors. Rather than following fixed developmental blueprints, type-specific dendritic architectures appear to largely emerge from the collective behaviors of self-organized local branch interactions^14,18,23–27^. According to these findings, transcription factors and extrinsic cues may achieve morphological reproducibility not by instructing final shape directly, but by constraining stochastic branch behaviors through cytoskeletal regulation. Yet, fundamental questions remain: Can a minimal set of stochastic rules explain how diverse dendritic architectures emerge? How do local cytoskeletal regulations shape stochastic dynamics into reproducible morphologies?

Addressing these questions has been challenging due to technical limitations in imaging neuronal morphogenesis *in vivo*, and to the lack of efficient image-processing tools and simple, universal computational models. In this study, we overcome these limitations to compare the early developmental dynamics of two morphologically distinct classes of *Drosophila* dendritic arborization (da) neurons. These mechanosensory neurons, which develop during late embryogenesis, are present in stereotypical locations in each embryonic segment (Figure 1A). They are accessible for live imaging and exhibit class-specific arbor patterns. They are classified into four morphological and functional classes (I–IV) of increasing complexity (Grueber et al., 2002). Class-I neurons, involved in proprioception, form simple comb-like arbors optimized for sensing epidermal deformation and coordinating muscle contractions (Figure 1A, D)^28–31^. In contrast, Class IV neurons which are polymodal nociceptors^32,33^, develop highly branched, space-filling arbors that tile the body wall, enabling efficient detection of localized noxious stimuli and rapid escape responses (Figure 1A, B)^34^.

**Figure 1.**
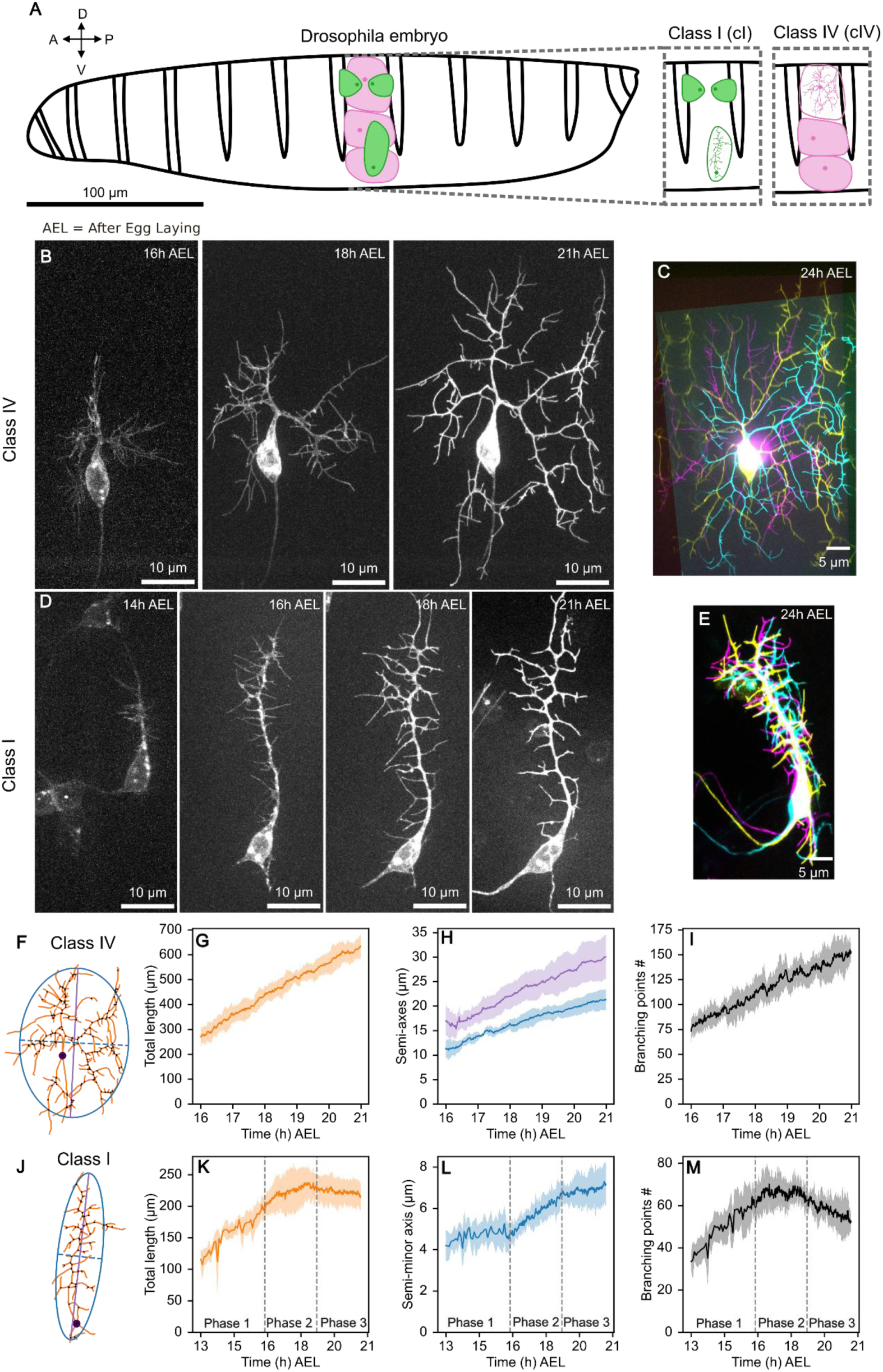
Distinct growth modes in Class I and Class IV neurons. (A) Stereotypical positions of *Drosophila* Class I (green) and Class IV (pink) dendritic arborization neurons in each embryonic/larval segment. This study focuses on ventral vpda Class I neurons (green circle) and dorsal ddaC (pink circle) Class IV neurons. (B-D) Time-lapse imaging of dendritic growth in Class IV neurons labelled by ppk-Td::GFP starting at 16h After Egg Laying (16h AEL) (B) and Class I neurons labelled by rluv-Gal4>UAS-CD4::NeonGreen (left, 14h AEL) or 221-Gal4>CD4::NeonGreen (16, 18 and 21AEL). Class-IV growth is stochastic, as illustrated by the overlay of three ddaC neurons in complementary colors (C). In Class I, growth of the primary branch is deterministic while higher-order branches grow stochastically, as illustrated by the overlay of 3 vpda neurons (E). (F-M) Growth morphometrics of Class IV (F-I) and Class IV (J-M) neurons illustrated in panels (F) and (J). The morphometric parameters used for comparison are: total arbor length (orange), vertical extension (purple line), lateral extension (blue dotted line) and number of branching points (black dots). Class-IV neurons grow linearly (G-I) whereas Class I neurons grow more slowly, in a stepwise manner (K-M). Three developmental phases are distinguished for Class I neurons. Phase 1: deterministic dorsal extension of the primary branch; Phase 2: stochastic growth and extension of higher order branches; Phase 3: refinement and stabilization of the dendritic arbor. Solid lines indicate the mean; shaded area indicates mean ± 1 SD. n=5 Class IV and n=6 (13h-16h) and n=7 (16h-21h) Class I neurons.

Class I and Class IV neurons have been extensively studied, leading to the identification of numerous genes required for their morphogenesis^35–42^. However, the dynamical principles shaping their characteristic architectures have only recently begun to be characterized with the help of computational modeling. Class IV neurons develop variable dendritic trees that seem to arise from stochastic branching, with no evidence of deterministic guidance (Figure 1B, C)^26,27^. In contrast, the reproducible comb-like shapes of Class I neurons emerge from the combined action of yet unidentified directional cues that guide the invariant dorsal extension of their primary branch, together with stochastic self-organized processes that shape the lateral extension of higher order branches (Figure 1D, E)^14,18^. Current computational models of Class I and Class IV morphogenesis are based on two or three discrete phases (growth, retraction and pause), rely on a large number of parameters and assume diffusive dendritic branch behavior. Although they successfully recapitulate local branch dynamics in both neuronal types, including stochastic growth and shrinkage, lateral branching, and contact-induced retraction (Figure 2A, G, and I), they do not yet reveal the fundamental dynamical principles underlying the emergence of their distinctive morphologies. In other words, how large-scale neuronal architecture is encoded in the stochastic dynamics of local branching processes remains unresolved.

**Figure 2:**
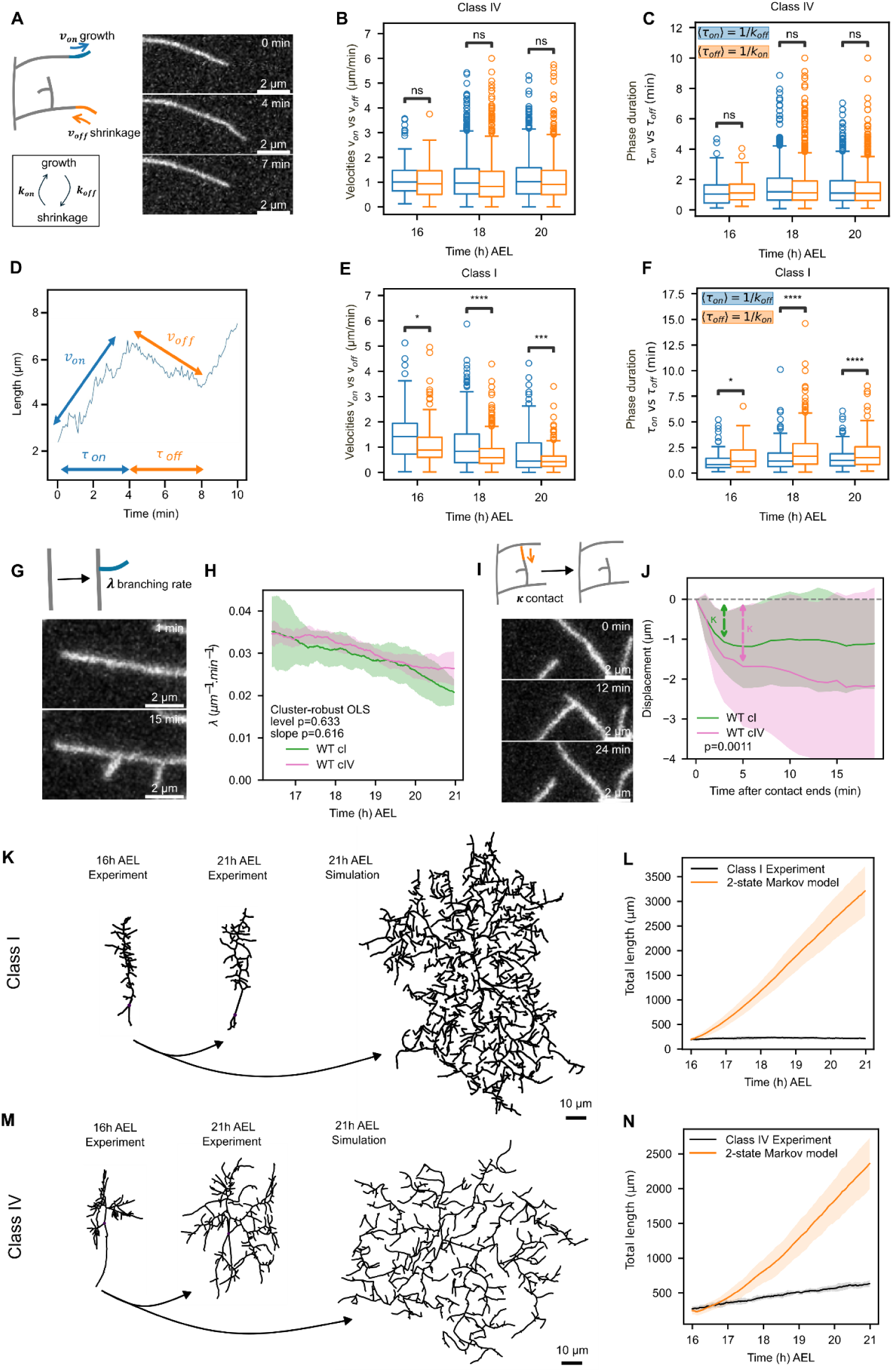
The two-state Markov model overestimates neuronal growth. (A-J) Measurements of local dynamical parameters. Both classes alternate between phases of growth and shrinkage (D), characterized by elongation and retraction velocities (v_on_, blue; v_off_ orange) and switching rates (k_on_=1/τ_off_, blue; k_off_=1/τ_on_, orange). Individual phases were automatically segmented using piecewise linear regression. (A) Schematic of growth and retraction phases and representative time-lapse imaging of branch dynamics in a Class IV (B, C) and a Class I neuron (B, C). Distributions of phase velocities (B,E) and durations (C,F) over developmental time. In Class IV neurons, mean growth and shrinkage velocities (⟨v_on_⟩ ∼ ⟨v_off_⟩) and durations (⟨τ_on_⟩ ∼ ⟨τ_off_⟩) are comparable; Two-sided Welch’s t-tests with Holm correction. Velocities: 16 h AEL, p = 0.211; 18 h AEL, p = 0.164.; 20 h AEL, p = 0.211. Durations: 16 h AEL, p = 1.00; 18 h AEL, p = 1.00; 20 h AEL, p = 1.00. (E, F) In Class I neurons, ⟨v_on_⟩ > ⟨v_off_⟩ (E) whereas ⟨τ_on_⟩ < ⟨τ_off_⟩ (F). Velocities: 16 h AEL, p = 2.06 × 10^-2; 18 h AEL, p = 1.83 × 10^-6; 20 h AEL, p = 1.35 × 10^-3. Durations: 16 h AEL, p = 5.23 × 10^-2. 18 h AEL, p = 2.56 × 10^-8. 20 h AEL, p = 4.42 × 10^-5. Quantifications were performed on high-temporal resolution movies (1 frame every 5s for 10 min) at 16, 18 and 20h AEL. Class-I: 21 acquisitions, 332 trajectories, 1393 phases. Class-IV: 16 acquisitions, 780 trajectories, 3448 phases. (G) Schematic and time-lapse imaging illustrating side-branching, quantified by the branching rate λ (number of new branches per unit time and per unit total dendritic length). (H) Branching rate λ (µm^-1^·min^-1^) over time for Class I (green, n= 7) and Class IV (pink, n = 5) neurons. λ decreases over time in both classes (Class I: p = 0.0416; Class IV: p = 0.019) but does not differ between them (level, p = 0.644; slope, p = 0.602). Statistics were computed using OLS with cluster-robust standard errors (clustered by acquisition, see Material and Methods). Curves were smoothed for visualization (Savitzky-Golay filter, see Material and Methods). Statistics were computed on unsmoothed data. (I) Time-lapse imaging of contact-induced branch retraction, quantified by κ (µm), defined as the mean retraction length following branch contact. (J) κ is greater in Class IV (–1.68 ± 0.17 µm, pink, n=79 branches) than in Class I neurons (–1.15 ± 0.0798 µm, green, n=127 branches). Welch’s t-test: p = 0.00582. (K-N) Neuronal growth simulations using the two-state Markov model. (K,M) Comparisons of Class I (K) and Class IV (M) neuronal morphologies at 21 AEL (middle panels) with simulated neurons using parameters extracted from *in vivo* data (right panels). Simulations were initialized from the corresponding neuron on the left (16 AEL). (L,N) Total arbor length measured in vivo (black) and in silico (orange) for Class I (L) and Class IV (N) neurons. The two-state Markov model substantially overestimates neuronal size and does not recapitulate their morphologies. In vivo, n = 7 (Class I), n = 5 (Class IV). In silico, n = 70 (Class I), n = 50 (Class IV). H, J, L, N. Data are mean ±SD.

Here, we aim to identify both the shared principles and the distinguishing features of dendrite morphogenesis in Class I and Class IV neuron. We compare the early shape changes of these neurons using high-resolution imaging, automated morphometric measurements, computational modeling and cytoskeletal perturbations. Despite sharing similar local branching rules, we show that the two classes follow distinct growth algorithms that cannot be explained by standard, diffusive models. By decomposing branch dynamics into short-term exploratory fluctuations and long-term subdiffusive behaviors, we reveal how actin and microtubules differentially shape arbor growth and morphology. A minimal model incorporating these dynamics accurately recapitulates class-specific growth trajectories and final architectures, providing a general framework linking local stochastic dynamics to global arbor expansion and reproducible dendritic morphology.

## Results

### The distinct morphodynamics of Class I and Class IV dendritic neurons

In light of the high complexity of the cellular and molecular mechanisms underlying neuronal morphogenesis, we first adopted a phenomenological approach to characterize the morphodynamics of Class I and Class IV neurons. Previous studies have mainly focused on larval stages, when the core morphology of these neurons is already established. To characterize and compare the gradual elaboration of Class I and Class IV dendritic arbors, we therefore tracked their early development during embryogenesis, using time-lapse movies acquired every minute for over 6 hours (Section 2 in Supplementary Material and Methods, Figure 1B,D, Movie S1). As shown previously, both neuron types follow similar local growth rules, including stochastic branch elongation and retraction, side branching and shrinkage induced by self-contact^14,18,26^. Yet, the two neuronal classes exhibit strikingly different overall tree growth dynamics. To quantify these differences, we developed image-analysis tools to characterize the global morphodynamics of dendritic trees (Section 3 in Supplementary Material and Methods, Tables 1 and 2). We measured each of the following parameters as a function of time: the total dendritic length (*L*), defined as the sum of all branch lengths, the semi-major and semi-minor axes of ellipses fitted to the dendritic trees, the number of branching points (*B*) and the branch density (Figure 1F-M and S1).

Class-IV neurons initiated dendritic outgrowth around 16 h <u>A</u>fter <u>E</u>gg <u>L</u>aying (AEL) (Figure 1B, 1F-I and S1G-L, Movie S1 and Table 1). Their total dendritic length *L* increased at a linear rate of 1.20 µm.min^-1^ (Figure 1G), accompanied by a linear increase in the number of branching points (0.25 per min, Figure 1I) (n=5 neurons). Arbor expansion was slightly anisotropic, as indicated by the semi-major and minor axes of the fitted ellipses, but their relative proportions remained constant over time (Figure 1H, S1J). Meanwhile, branch density gradually decreased (Figure S1L). In contrast, Class I neurons developed more slowly, through three sequential linear phases (Figure 1D, 1J-M and S1A-F, Movies S1 and S2, Table 2): During the first phase (13–16 h AEL), the primary branch elongated dorsally at a rate of 0.41 µm.min^-1^, accompanied by a fast increase in the number of branching points (0.14 per min) but minimal lateral expansion (phase 1, n = 7). During the second phase (16–19 h AEL), higher-order branches extended laterally, while the overall growth rate and arbor density progressively declined, and the number of branching points plateaued (phase 2, n = 7). Finally, during the third phase (19–24 h AEL), arbor development slowed further, with a slight reduction in total dendritic length, modest lateral spread and a decrease in both branching points and density (phase 3, n = 7). Thus, Class IV neurons grow rapidly and continuously, whereas Class I neurons develop in a slower, stepwise fashion. This is consistent with their reduced size and increased anisotropy relative to Class IV neurons.

### A two-state Markov model fails to account for dendritic morphogenesis

We then sought to understand how the divergent growth of Class I and Class IV neurons emerges from differences in the local stochastic dynamics of their branching processes. To address this question, we first defined a minimal parametrization of these dynamics, inspired and constrained by the observed phenomenology. We then used computational modelling to simulate dendritic morphogenesis and compared the simulations with live-imaging data, to determine which dynamical model best captures morphogenesis of both neuronal types.

We initially considered a previously developed two-state Markov dynamical model that describes the stochastic behavior of second-and higher order-branches of Class I neurons (Section 8.2.1 in Supplementary Material and Methods)^14^. In this two-state model, branches alternate between phases of growth and retraction with characteristic velocities v_on_ and v_off_ (Figure 2A, D). Transitions between these states follow a continuous-time Markov chain, governed by switching rates k_on_ (from retraction to growth, min^-1^) and k_off_ (from growth to retraction, min^-1^) (Figure 2A). New branches emerge laterally from existing ones with rate λ (µm^-1^.min^-1^) (Figure 2G), and branches retract upon contact with neighboring branches with an amplitude κ (µm) (Figure 2I). All parameters are assumed to be statistically independent and the memoryless nature of the Markov chain implies that successive growth and shrinkage phases are independent, resulting in diffusive branch dynamics. A limitation of the original implementation of this model in Class I neurons is that parameter values were not measured *in vivo*. In the present study, we directly quantified all local dynamical parameters *in vivo*, reasoning that class-specific differences in these parameters should, when implemented into the model *in silico*, quantitatively account for the neuronal morphodynamics observed experimentally.

To quantify branch-tip velocities and switching rates, we analyzed non-contacting branches in high-temporal-resolution movies (one frame every 5s for 10min) at 16, 18, and 20 h AEL (Movies S3 and S4). Segmentation of 1112 distinct branches from 21 Class I (n= 332 branches) and 16 Class IV (n=780 branches) neurons yielded independent time evolutions of branch length, which we refer to as branch trajectories (Section 2.4 in Supplementary Material and Methods, Figure 2D, S2A-F Table 3). These trajectories were further segmented into individual steps revealing growth (blue) and shrinkage (orange) events characterized by specific velocities (v_on_, v_off_) and durations (τ_on_ and τ_off_) (Figure 2D and S2G,J, Section 7 in Supplementary Material and Methods). Population statistics revealed exponential distributions across these metrics (Figure S3), from which we computed the mean values v_on_, v_off_, k_on_ (=1/<τ_off_>) and k_off_ (=1/<τ_on_>) as a function of time for both neuronal classes (Figure 2B-F, Table 4). These analyses revealed some differences between the two neuronal classes. In Class IV neurons, elongation and retraction occurred at similar speeds (v_on_ ∼ v_off_ ∼1µm.min^-1^) and switching rates (k_on_ ∼ k_off_ ∼0,71min^-1^) and all parameters remained unchanged over time (Tables 4 and 5, Figure 2B,C). In Class I neurons, however, elongation speed consistently exceeded retraction (v_on_ > v_off_), and both slowed over time, but branch tips were more likely to switch from growth to retraction (k_off_ > k_on_) (Tables 4 and 5, Figure 2E,F). In early stages, Class I branches elongated as fast and as often as Class IV branches, but from 18h onward they retracted more rapidly and more frequently (Figure S2H-L, Table 5).

We next measured the branching rate *λ*—defined as the number of new branches formed per unit of branch length and time —using time-lapse movies acquired every minute over a 5 hours period (Section 2.3 in Supplementary Material and Methods, Movie S1, Figure 2G). *λ* was similar between Class I and Class IV neurons (0.036 µm^-1^.min^-1^ at 16 h AEL) and decreased gradually over time (Figure 2H, Tables 6 and 7). From the same movies, we also extracted the mean length κ of branch retraction after branch contact (Figure 2I) and found that it was slightly larger and persisted longer in Class IV neurons than Class I neurons (1.15 µm for Class I and 1.68 µm for Class IV, Figure 2J, Tables 8 and 9).

At first glance, the measured differences in local branch dynamics between Class I and Class IV neurons did not provide an intuitive explanation for their distinct morphologies. Some parameters, such as λ, remained consistently similar between the 2 classes over time, whereas k_on_, k_off_, v_on_ and v_off_ differed by up to 50% depending on the developmental stage (Figure S2, Table 4). However, in both cases, the mean branch velocity computed from the respective values of v_on_, v_off_, k_on_ and k_off_ (C= k_on_ /(k_on_ + k_off_).v_on_ – k_off_ /(k_on_ + k_off_).v_off_) did not differ significantly from zero in either neuronal type (drift in Table 4), indicating no net directional motion at the level of individual branches. Nevertheless, dendritic arbor growth emerges from the cumulative effect of stochastic elongation and branching events over long timescales. To determine whether these modest differences in local dynamical parameters could nonetheless generate the observed morphological differences between Class I and Class IV neurons, we used computational modeling. We simulated dendritic growth using the 2-state Markov model parametrized by *in vivo* measurements for each neuronal class. Model predictions were compared to live-imaging data by systematically analyzing the time evolution of the same morphometric parameters computed from the experiments (Figure 1F, J).

The initial condition of our simulations was the skeleton of one neuron from each class extracted at the start of live-imaging acquisitions (10 simulations per neuron, 7 Class I neurons, 5 Class IV neurons; Movie S5 and S6). Strikingly, the simulations greatly overestimated neuronal growth in both neuronal classes: arbor total length, lateral expansion (semi-minor axis), and the number of branching points all exceeded experimental values, by 2-4-fold higher in Class IV and 6-16-fold in Class I neurons (Figure 2K-N and S4A-H). Moreover, there was no morphological distinction between simulated Class I and Class IV neurons (compare Figure 2K and M) whereas the *in vivo* data show clear differences in size and shape. Thus, despite the measured class-specific differences in local branch-tip dynamics, the two-state Markov model failed to account for the observed morphologies, indicating that some of its underlying assumptions are incorrect.

### Velocity-duration coupling underlies the short-term branch diffusivity of branches

The first key assumption of the two-state model is that the dynamical variables are statistically independent, in particular the durations (τ_on_ and τ_off_) and velocities (v_on_ and v_off_) of individual growth and shrinkage events. To test this assumption, we analyzed individual growth and shrinkage steps in detail. We found that *in vivo*, step velocity and duration were correlated in both neuronal classes: slower steps tended to last longer whereas faster steps were shorter-lived (Figure 3A, D and S5A-F, Table 10). This finding indicates that the two-state model overestimates the occurrence of long-lasting, high-velocity events, artificially amplifying branch-length fluctuations in simulations. This may explain the excessive neuronal growth and size predicted by this model compared with experimental data.

**Figure 3.**
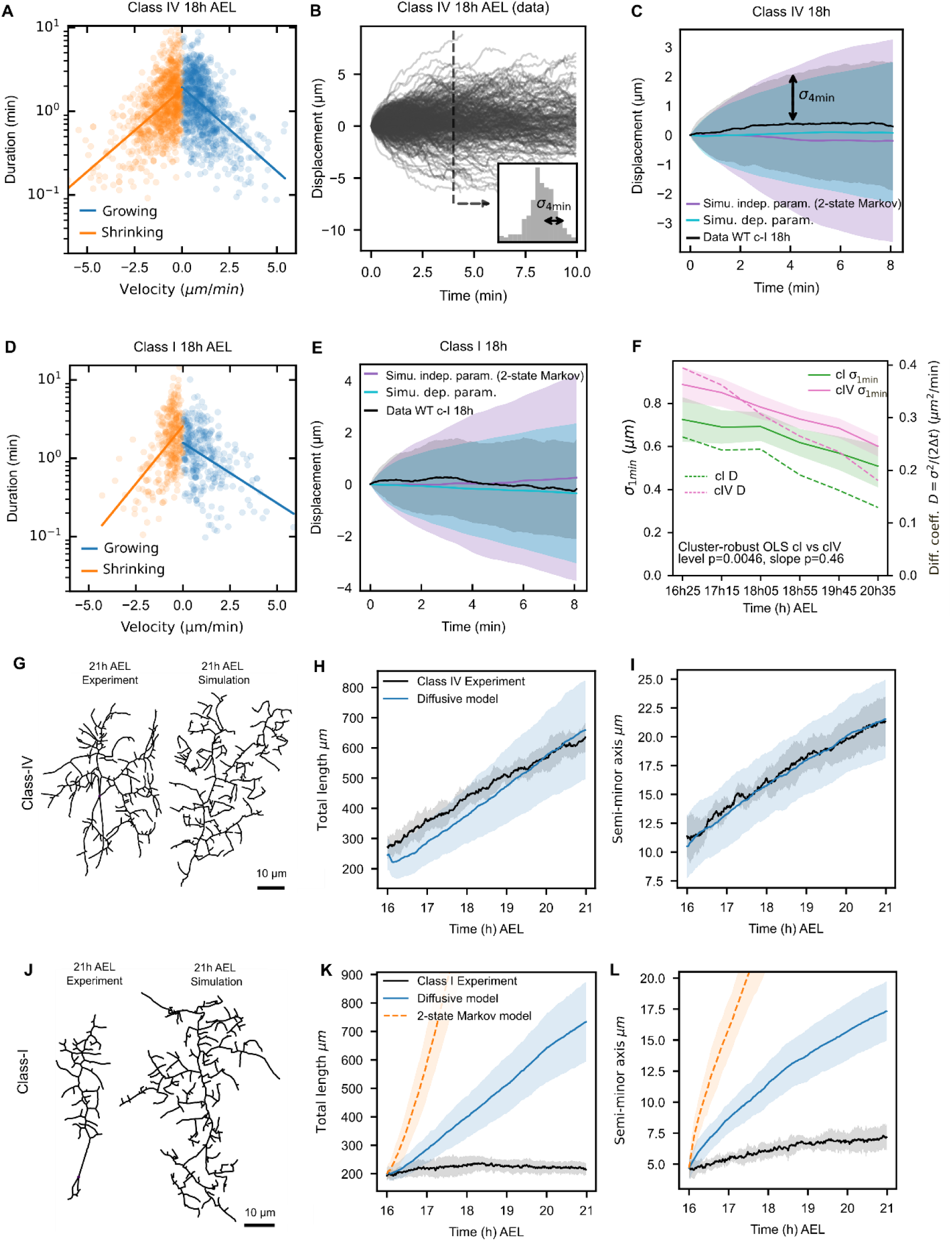
A coarse-grained model resolves diffusion overestimation. (A,D). Joint distributions of phase velocity and phase duration at 18 h AEL. Each point represents a single phase (growth in blue, shrinkage in orange). Solid lines indicate linear regressions between velocity and log(duration), revealing coupling between the two variables. (A) Class IV neurons: growth slope = –0.46 (p = 2.8e-60), shrinkage slope = –0.45(p = 5e-64). (D). Class I neurons: growth slope = –0.36 (p = 5.1e-14), shrinkage slope = –0.68 (p = 8.8e-23). (B) Displacement of individual branch tips, aligned to 0 µm at t = 0 min. Inset: distribution of displacements at 4 min with corresponding standard deviation (σ_4min_). (C,E) Comparisons of mean displacement of branch-tip trajectories from experimental data (black), simulations from the 2-state Markov model assuming independence between velocity and duration (purple), and a model incorporating velocity-duration coupling (cyan) in Class IV (C) and Class I (E) neurons. Diffusion coefficients D are reported with bootstrap 95% confidence intervals. P values are from two sided bootstrap tests on ΔD (B = 200 resamples). In Class IV neurons (C), the independence model overestimates diffusion (data: D = 0.30 µm^2^.min^-1^, simulations: D = 0.75 µm^2^.min^-1^, p < 0.01), whereas the coupling model is not significantly different from experimental data (D = 0.36 µm^2^.min^-1^, p = 0.14). In Class I neurons (E), the independence model also overestimates diffusion (data: D = 0.20 µm^2^.min^-1^, simulations: D = 1.0 µm^2^.min^-1^, p < 0.01); incorporating velocity-duration coupling reduces this overestimation, but D remains different from experimental data (D = 0.48 µm^2^.min^-1^, p < 0.01). Analysis is performed on trajectories that last at least 8 min: n=170 for class IV and n=67 for class I. (F). Standard deviation of 1 min displacement increments (σ_1min_) over time. σ_1min_ decreases in both classes (Class I, green, p = 0.028; Class IV, magenta, p = 0.0033). was significantly higher in Class IV than in Class I neurons at 16h25 AEL (cluster-robust OLS: level, p = 0.00465), whereas the temporal slopes did not differ significantly between classes (p = 0.46). Dashed lines indicate diffusion estimates derived from σ_1min_. (G-L) Neuronal growth simulations using the coarse-grained diffusive model incorporating velocity-duration coupling. Comparisons of Class I (G) and Class IV (J) neuronal morphologies at 21h AEL (left panels) with simulated neurons using parameters extracted from in vivo data (right panels). Simulations were initialized from the corresponding neurons shown in Figure 2K (Class I) and 2M (Class IV). (H,I,K,L) Morphometric parameters measured in vivo (black) and in silico (blue). The coarse-grained diffusive model accurately recapitulates Class IV neurons growth: overall morphology is well reproduced (G), and total arbor length (H) and semi-minor axis (I) fall within the correct range. In contrast, for Class I neurons, final morphology is too large (J) and morphometric parameters remain overestimated (K,L). *In vivo*: n = 7 (Class I); n = 5 (Class IV). In silico: n = 70 (Class I) and n = 50 (Class IV). C,E,F,H,I,K,L. Data are mean ± SD.

To test this further, we analyzed and compared the behavior of individual branch trajectories in experimental and simulated datasets. In a diffusive process such as the two-state Markov model, random fluctuations in branch length cause branch-tip displacement to spread over time without directional bias. This behavior can be described by an effective diffusion coefficient *D,* defined as the proportionality coefficient between branch displacement variance and time. We first superimposed 170 branch trajectories extracted from 10 min time-lapse recordings acquired every 5 seconds from 10 Class IV neurons at 18h AEL (Figure 3B and Table 11). The dispersion of these trajectories (grey envelope in Figure 3C) is reflected by the time evolution of the standard deviation σ_*t*_ of the displacement distribution (e.g., σ_4min_, Figure 3B). The diffusion coefficient of these branches D_data_ was estimated from the linear fit of the Mean Square Displacement (MSD) E[Δx(t)^2^] as a function of time, where Δx is the displacement of a branch over the time interval t (E[Δx(t)^2^]=2Dt, (Section 7.6 of Supplementary Material and Methods). This yielded a value of 0.30µm^2^.min^-1^ (Figure 3C). Simulations of independent branch trajectories assuming independent variables (two-state Markov model, section 7 in Supplementary Material and Methods) produced a much broader displacement distribution (purple envelope in Figure 3C) and a substantially higher diffusion coefficient (D_ind_=0.75µm^2^.min^-1^). When the experimentally observed dependencies between τ_on_, τ_off_, v_on_ and v_off_ were used in the simulations of individual branches, the distribution of branch displacements became narrower and closer to *in vivo* measurements (blue envelope Figure 3C). The resulting diffusion coefficient (D_dep_=0.36µm^2^.min^-1^) was also closer to experimental value. In Class I neurons, 67 trajectories from 8 neurons were analyzed (Figure 3E and Table 11). Branch trajectories were much less diffusive in experimental data (D_data_ = 0.2µm^2^.min^-1^) than in simulations assuming independent variables (D_ind_=1.0µm^2^.min^-1^). As in Class IV neurons, introducing the observed dependencies reduced diffusivity and the extent of branch displacement (D_dep_*=*0.48µm^2^.min^-1^, p<0.01), bringing simulations closer to the experimental value (Figure 3E). Similar results were obtained for trajectories extracted at 16 and 20 h AEL in both neuronal classes (Figure S5G-L). These results demonstrate that the experimentally observed coupling between velocity and step duration is essential for accurately capturing the diffusivity of Class I and Class IV individual branches in simulations.

Building on this finding, we next incorporated velocity–duration coupling into whole-neuron simulations, enabling us to capture how branch dynamics shape arbor geometry. We tested whether this improves predictions of neuronal size and shape compared with the two-state Markov model. To this end, we developed a coarse-grained diffusive model (section 8.2.2 in Supplementary Material and Methods), in which the size of individual steps, Δx, was statistically derived from the standard deviation σ of the branch displacement distribution measured from experimental data. σ naturally captures the *in vivo* coupling between step duration and velocity and can be used to estimate the diffusion coefficient D (D= σ ^2^/2Δt, section 7.6 of Supplementary Material and Methods). To simulate neuronal growth over long timescales, we estimated σ from 5h time-lapse recordings acquired a 1min intervals, which we also used to measure branching and contact-induced retraction parameters λ and κ (Movie S1, Figure 2G,I). In Class IV neurons, σ was 22% higher than in Class I neurons (level at 16h25: p=0.00465, Table 13) and decreased over time in both classes (Figure 3F, Table 12) at a similar rate (p = 0.46, Table 13). Remarkably, introducing velocity-duration coupling strongly reduced the total arbor length, semi-minor axis and number of branching points in simulated Class IV neurons compared to two-state model with independent variables (compare Figures 3G-I and S4M-P to Figures 2M-N and S4E-H, Movie S7). Including the drift in the distribution of branch length variations Δx did not improve the model’s fit to the data (Figure S6A,B,D-G, section 7.7 in Supplementary Material and Methods). In contrast, for Class I neurons, the model still overestimated total arbor length, the semi-minor axis and the number of branching points of ∼3-fold, producing abnormally shaped neurons compared with experimental data (Figure 3J-L and S4I-L, Movie S8). This overestimation persisted irrespective of whether the drift in the distribution of Δx was included or not (Figure S6A,C,H-K). Thus, correcting the scale of branch exploration through velocity-duration coupling was sufficient to recapitulate Class IV morphology, but not that of Class I neurons.

### Long term anomalous branch diffusion recapitulates class-specific divergent morphologies

The persistent overestimation of Class I growth metrics pointed to a second potential limitation of our models: successive steps of growth and shrinkage may not be independent in these neurons as assumed so far. Consequently, branch fluctuations may deviate from diffusive dynamics over long timescales. To test this, we quantified the long-term (>1 h) variability of branch trajectories in Class I and Class IV neurons, using the 1 min resolution time-lapse recordings over 5 hours and computed the MSD (Movie S1). Time evolution of branch length trajectories in 7 Class I and 5 Class IV neurons revealed more constrained branch dynamics in Class I compared with Class IV neurons (Figure 4A, Table 14). Under normal diffusion, MSD increases linearly with time (MSD(t) ∝ t). Consistent with this, branch fluctuations in Class IV neurons were diffusive, with MSD(t) ∝ t^α^ and α=1.00±0.21 (Figure 4B). In contrast, Class I neurons exhibited strongly subdiffusive behavior, with α=0.45±0.14 (Figure 4B). The difference in the anomalous diffusion exponent α between the two classes was highly significant (Figure 4C, p=4.3 x10^-5^). This pronounced subdiffusive behavior in Class I neurons likely explains why diffusive models with independent (two-state Markov) or coupled (coarse-grained) variables systematically overestimated Class I arbor expansion. Notably, α is the main parameter that distinguishes Class I and Class IV neurons.

**Figure 4.**
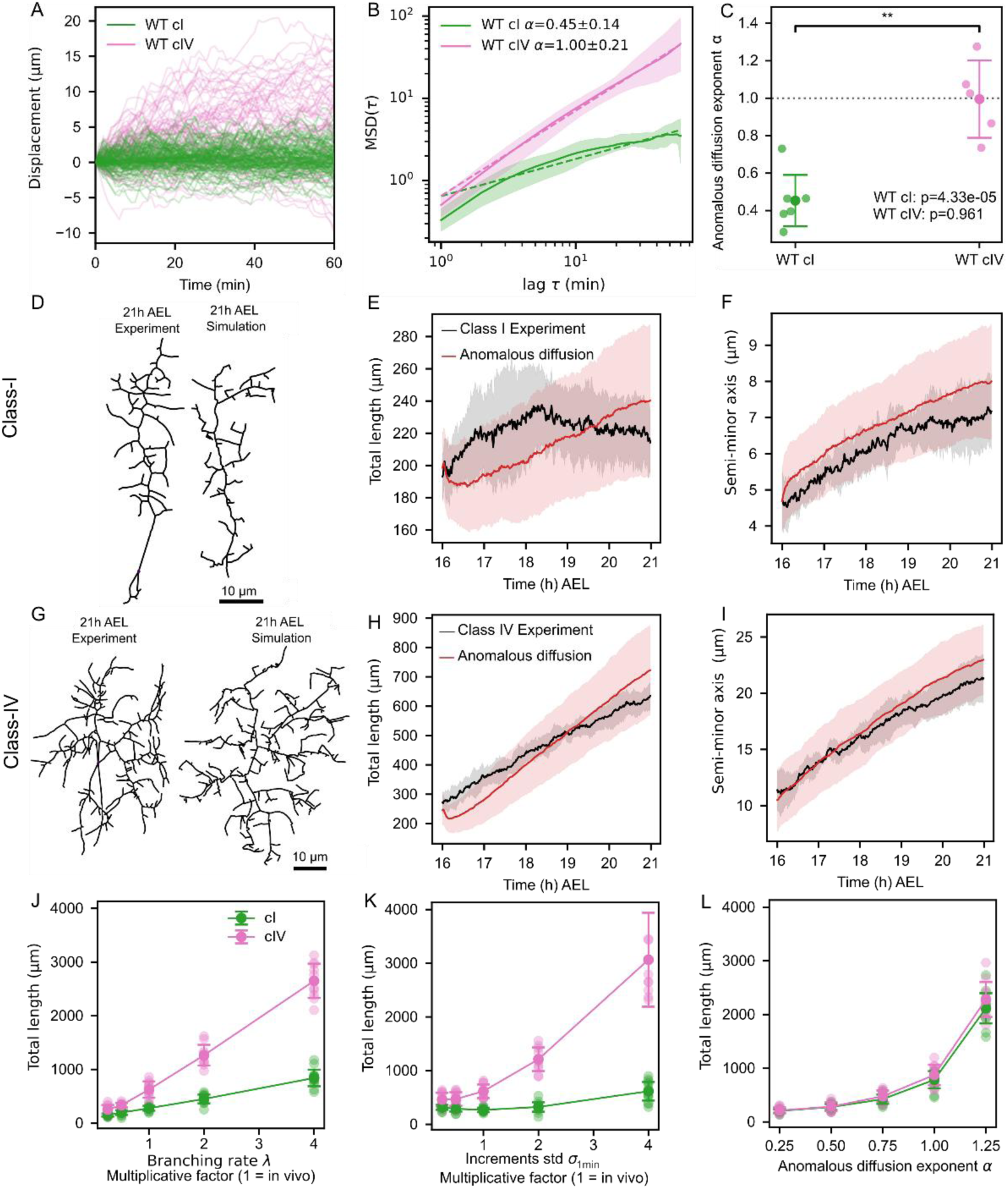
A parsimonious model incorporating the anomalous diffusion coefficient α captures the growth of Class I and Class IV neurons. (A) Displacement of individual branch tips, aligned to 0 µm at t = 0 min, in Class I (green) and Class IV (pink) neurons. (B). Mean squared displacement (MSD) for Class I (n = 7) and Class IV (n = 5) neurons. Solid line indicates the mean; shaded regions represent the range (min-max) across neurons. Dashed lines correspond to the fits used to estimate the anomalous exponent α, defined as the slope of the linear fit of log(MSD) versus log(t). (C) The anomalous exponent α differs between Class I and Class IV (two-sided Welch’s t-test, p = 0.00171). In Class I, α differs from 1 (two-sided one-sample t-test, p = 4.33e-5), indicating subdiffusive behavior. Class-IV neurons are diffusive (α=1, p = 0.961). (D-I) Neuronal growth simulations using the anomalous diffusion model. Comparisons of Class I (D) and Class IV (G) neuronal morphologies at 21h AEL (left panels) with simulated neurons using parameters extracted from *in vivo* data (right panels). Simulations were initialized from the corresponding neurons shown in Figure 2K (Class I) and 2M (Class IV). (E,F,H,J) Morphometric parameters measured *in vivo* (black) and *in silico* (red). The anomalous diffusion model accurately recapitulates Class I and Class IV neurons growth: overall morphologies are well reproduced (compare left and right panels in D,G) and total arbor length (E,H) and semi-minor axis (F,I) fall within the correct range. Note that for Class IV, the anomalous diffusion model reduces to the diffusive model since α = 1 in these neurons. In vivo: n = 7 (Class I); n = 5 (Class IV). In silico: n = 70 (Class I) and n = 50 (Class IV). Data show mean ± SD. (J-L). Total arbor length in simulations while varying one parameter of the anomalous diffusion model at a time, with all other parameters fixed to class-specific measured values. For λ and σ, the x-axis represents the multiplicative factor relative to the reference value (x=1). Setting λ (J) or σ (K) to identical values does not change baseline morphological differences between Class I and Class IV, and increasing these parameters further amplifies them. In contrast, imposing the same α in both classes suppresses these differences (L), indicating that α is the primary parameter distinguishing Class I and Class IV morphologies.

Based on these findings, we developed a parsimonious dynamical model incorporating branch subdiffusive behaviors (Section 8.2.3 in Supplementary Material and Methods). This model, hereafter referred as the anomalous diffusion model, is based on only four parameters (λ, κ, σ, α). Branching and contact-induced retraction rules λ and κ are retained, while contact-free branch tip dynamics are decomposed into short– and long-term components: short-term fluctuations are captured by σ (standard deviation of 1 min branch increments), and the long-term behavior is governed by the anomalous diffusion exponent α, which sets the temporal scaling of MSD (MSD(t) ∝ t^α^). Setting α = 1 reduces to the coarse-grained diffusive model assessed earlier (Section 8.2.2 in Supplementary Material and Methods).

Simulations of neurons using this minimal model and the experimentally measured parameters reproduced most experimentally observed features of growth dynamics in both classes of neurons (Figure 4D-I and S4Q-X). In Class I neurons, arbor density and the number of branching points were slightly underestimated but remained within the correct quantitative range, whereas total arbor length and lateral spread were consistent with experimental values (Figure 4D-F, Figure S4Q-T and Movie S9). In Class IV neurons, all morphometric features closely matched experimental observations (Figure 4G-I, Figure S4U-X and Movie S10). Together, these findings demonstrate that incorporating anomalous diffusion through the exponent α is essential to accurately recapitulate neuronal morphodynamics, and class specific differences in neuronal shape.

We next assessed how individual model parameters influence Class I and Class IV growth dynamics. To this end, we performed a series of simulations in which one parameter was set to the same value and systematically varied in both neuronal classes, while all other parameters were fixed at their empirically measured, class-specific values. Morphometric features were extracted at the end of each simulation for comparison. For the branching rate (λ), the standard deviation of short-term branch-length changes (σ) and the post-contact branch retraction amplitude (κ), a reference value (x=1 in Figures 4J,K, and S7) corresponding to the mean of in vivo Class I and Class IV values was scaled by a multiplicative factor ranging from 0.25 to 4. For the anomalous diffusion exponent α, assigned values range from 0.25 to 1.25, with α>1 corresponding to superdiffusive behavior. Increasing λ, σ or α significantly increased total arbor length (Figure 4J-L), semi-minor axis (Figure S7E-H) and branching point number (Figure S7I-L) in both neuronal classes. In contrast, variation of κ selectively affected arbor density (Figure S7B,F,J,N). Arbor density increased with λ and α but decreased with σ and κ (Figure S7M-P). The aspect ratio, largely constrained by initial morphology, converged toward similar values at high λ, σ, or α (Figure S7E-H). Strikingly, assigning identical values of λ and σ in simulated Class I and Class IV neurons and progressively increasing them amplified baseline morphological differences such as arbor length (Figure 4J,K). In contrast, assigning identical values of α markedly suppressed class-specific differences, even at low α (Figure 4L). These results indicate that the branching rate (λ) and the amplitude of short-term branch-length fluctuations (captured by σ) scale growth and can accentuate initial differences, whereas the anomalous diffusion exponent (α) acts as the primary determinant of class-specific dendritic architecture and neuronal morphology.

Our model predicts that, at the macroscopic scale, dendritic arbor growth proceeds as a propagating bulk with an approximately constant internal density behind an invasion front (Supplementary Information Figure SI4A). To determine how the local stochastic growth rules governed by λ, σ, α and κ set the pace of global arbor extension, we developed a theoretical framework that yields a mathematical expression for the speed (c) of the neuron’s invasion front (see Supplementary Information). Previous work has shown that tip-splitting branching random walks generate Fisher-KPP-type invasion fronts^43,44^, a framework later applied to branching morphogenesis in the mammary gland^3^. However, our system differs in two key aspects: new tips are born through side branching along pre-existing dendrites rather than being localized at the invasion front, and branch-tip motion can be subdiffusive rather than diffusive. These differences inherently alter front dynamics. In the diffusive limit with tip splitting, the classical Fisher-KPP model yields the mean-field front speed (Supplementary Information):

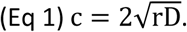

However, introducing side branching modifies the mean-field front speed to:

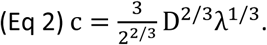

More generally, for subdiffusive branch-tip dynamics with anomalous exponent α, we obtained

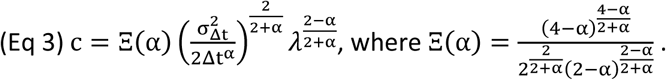

Importantly, sensitivity analysis confirmed a clear parameter hierarchy: front speed depends more strongly on the anomalous diffusion α than on σ, which reflects short term dynamics, and more on σ than on the branching rate λ in the relevant parameter range. Two-dimensional neuron simulations showed lower front speeds (Supplementary Information Figure SI4B) and slightly larger fitted exponents for σ and λ than the one-dimensional theory (Supplementary Information Figure SI4C) but preserved the parameter hierarchy. This provides an explanation for why long-timescale branch dynamics, governed by α disproportionately shape arbor expansion and class-specific morphology (Supplementary Information).

### Microtubules tune branch long term anomalous diffusion and arbor expansion

The branching rate (λ) is similar between Class I and Class IV neurons and short-term dynamics (σ) are modestly higher in Class IV neurons. However, long-term dynamics, quantified by α, differ markedly. These differences prompted us to investigate the cellular mechanisms underlying branch dynamics and their associated parameters. Since dendritic branch behavior depends on cytoskeleton activity, we examined the cytoskeletal basis of the class-specific differences, focusing on the respective contributions of actin filaments and microtubules. We generated fly lines enabling simultaneous *in vivo* imaging of F-actin (m-Scarlet-I::UtABD), microtubules (mNeonGreen::αTub) and the plasma membrane (HALO_7_::CAAX, see Materials and Methods; Figure S8). In both neuronal classes, actin was enriched in branch tips (Figure 5A,B and S9A’, B’, C’), while microtubules were concentrated in the proximal regions of dendrites and gradually invaded actin-rich branches as they stabilized (Figure 5A’,B’ and S9A”,B”,C”) with the exception of the Class I primary branch, whose invariant vertical elongation was associated with high microtubule levels from the earliest stages (Movie S11 and S12).

**Figure 5.**
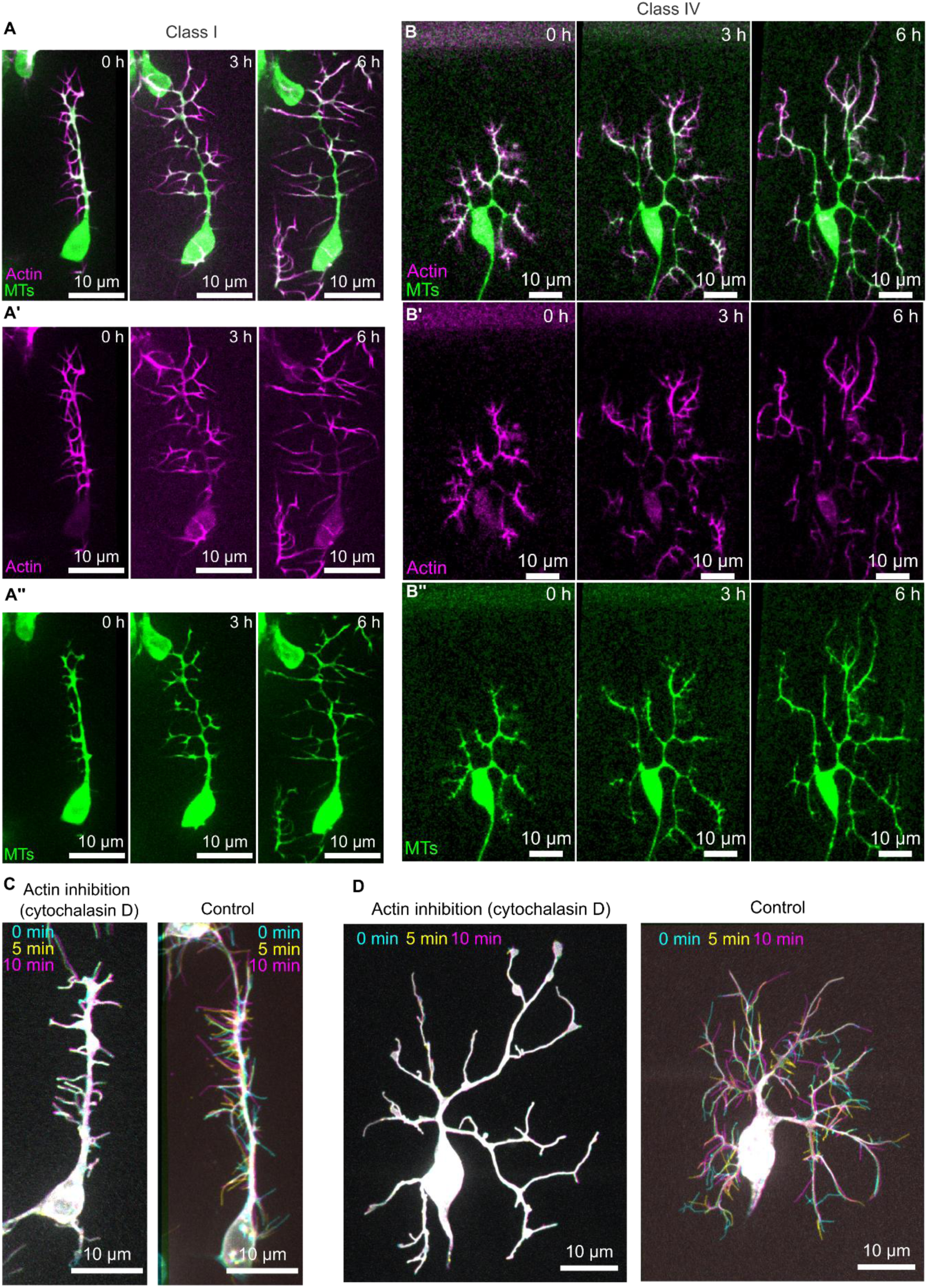
Cytoskeleton organization and actin perturbations in Class I and Class IV neurons. (A,B) Time-lapse imaging of the cytoskeleton in Class I (A-A”) and Class IV (B-B”) neurons during embryogenesis. Actin (mScarlet-I::UtABD) is shown in magenta and tubulin (*Neongreen::*α*-tub*) in green. In both classes, actin is enriched at dendritic tips, (A’, B’) while tubulin is mainly enriched in central regions and progressively invades the periphery of the dendritic arbor (A”, B”). (C,D) Actin inhibition in Class I (C) and Class IV (D) neurons via cytochalasin-D injection, with matched controls. Left panels show cytochalasin-D-treated neurons; right panels show controls. Regions with no displacement appear in white. Actin inhibition suppresses terminal branch dynamics in both neuronal classes.

To assess the respective roles of actin and microtubules during early development of Class I and Class IV neurons we perturbed cytoskeletal dynamics. Treatment with the actin polymerization inhibitor Cytochalasin-D completely abolished tip motility and branching in both neuronal classes, effectively freezing arbor development (Figure 5C,D, Movie S13 and S14). Consistent with this, the short-term dynamics captured by σ were reduced to near zero (not shown), indicating that actin controls both branch initiation and the short-term fluctuations that drive arbor expansion. In contrast, partial microtubules depolymerization (Colcemid + Vinblastine) in Class IV neurons, strongly reduced overall dendritic growth while leaving short term fluctuations largely unaffected (Figure 6A-H, Movie S15). Under these conditions, total arbor length increased by only ∼180 µm in 5 h (instead of ∼380 µm, Figure 6C) and lateral extension was limited to ∼2.5-fold increase (instead of ∼10-fold, Figure 6D), approaching *wild-type* Class I values (Figure 1L). The number of branching points was reduced (Figure 6E) whereas the branching rate λ showed only a modest decrease in overall levels (Figure 6F, Tables 6 and 7) and the standard deviation of short-term fluctuations σ remained unchanged (Figure 6G, Tables 12 and 13). In contrast, the anomalous diffusion exponent α was significantly decreased compared to *wild-type* (α=0.62, p=0.022, Figure 6H), and became non-significantly different from that of *wild-type* Class I neurons (α=0.45, P=0.066). Similar effects were observed in Class I neurons (Figure S10), although they were stage-dependent: early perturbations impaired primary branch elongation and lateral branch initiation (Figure S10A,A’), whereas later treatment preserved short-term fluctuations but markedly reduced final arbor size (Figure S10B,B’, Movies S16 and S17). Together, these results reveal distinct roles for actin and microtubules in the control of branch dynamics. Actin, enriched at the tips of growing branches, drives exploratory short-term fluctuations (σ), whereas microtubules, localized within the dendritic shaft, promote branch stabilization and global arbor expansion, by regulating the long-term diffusive behavior of branches.

**Figure 6.**
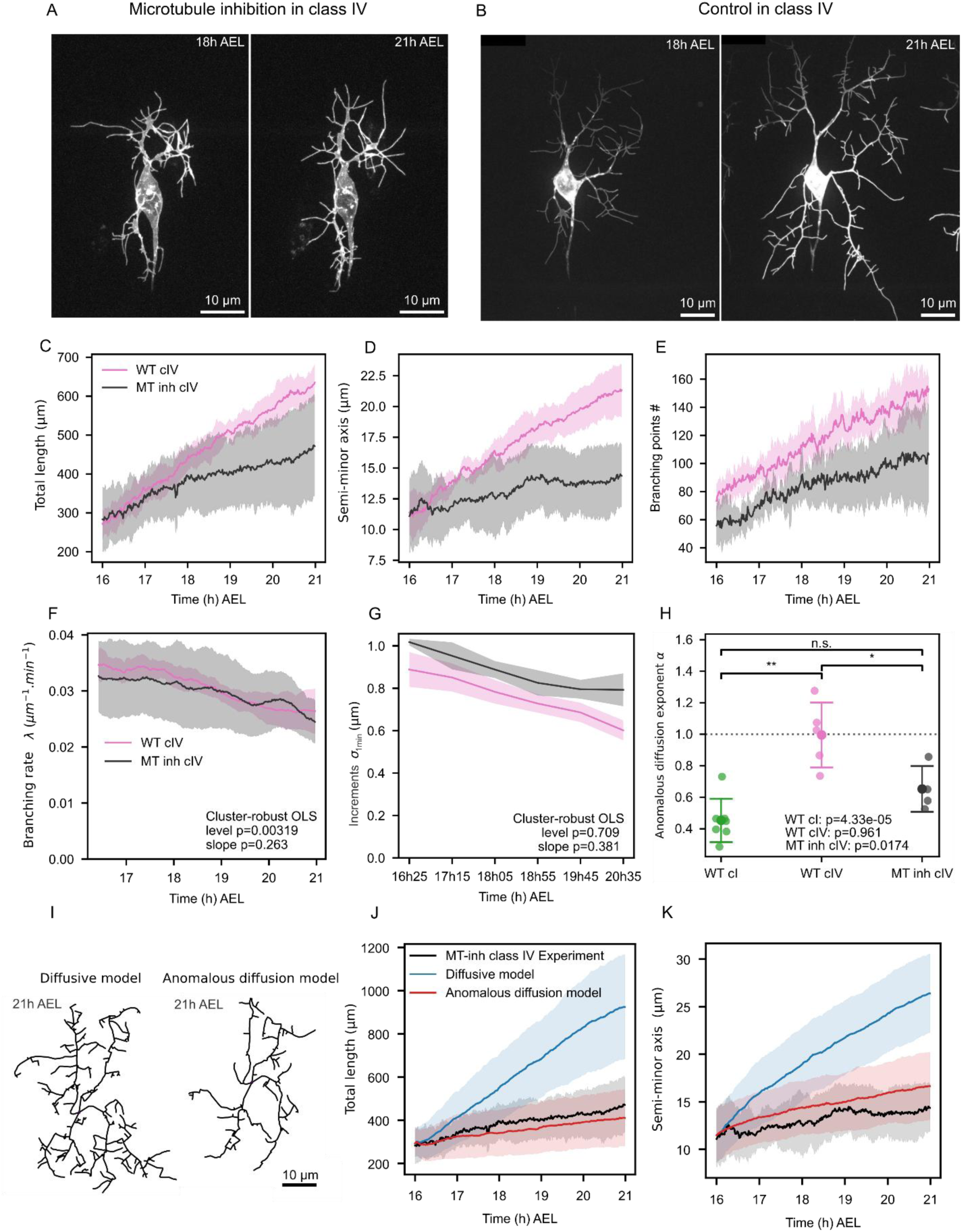
Microtubules inhibition in Class IV neurons. (A,B). Microtubules inhibition in Class IV neurons. Time-lapse imaging of a Class IV neuron following injection of Colcemid and Vinblastine (A) or water (B). Microtubules inhibition strongly impairs Class IV neurons dendritic growth. (C-E) Morphometric parameters of *wild-type* Class IV neurons (WT cIV, pink) and microtubule-inhibited Class IV neurons (MT inh cIV, black). Microtubules inhibition reduces total arbor length (C), lateral extension (D), and the number of branching points (E). (F-H). Evolution of dynamical parameters of the anomalous diffusion model in microtubule (MT)-inhibited Class IV neurons. (F) Branching rate λ in *wild-type* (WT) Class IV neurons (pink, n= 5) and MT-inhibited Class IV neurons (black, n= 4). λ decreases over time in both conditions (WT cIV: p = 0.019, MT-inhibited cIV: p = 0.0249; cluster-robust OLS, Table 6). MT inhibition produces a statistically significant but modest reduction in the overall λ level relative to WT Class IV neurons (p = 7.09 × 10-5), while slopes do not differ significantly (p = 0.308, cluster-robust OLS, Table 7). Curves were smoothed for visualization. Statistics were computed on unsmoothed data. (G) Standard deviation of 1 min displacement increments (σ =σ_1min_) over time. σ_1min_ decreases in MT-inhibited neurons (p = 0.0028, Tables 12 and 13), but does not different significantly from WT in either level (p = 0.71) or slope, (p = 0.38, cluster robust OLS: level). (H) The anomalous diffusion exponent α significantly deviates from 1 in MT-inhibited Class IV neurons (two-sided one-sample t-test, p = 0.0174). α differs significantly between MT-inhibited and WT Class IV neurons (Welch’s t-test, p = 0.0226), but is not significantly different from WT Class I neurons (Welch’s t-test, p = 0.0666). (I-K) Comparison of neuronal growth simulations of MT-inhibited Class IV neurons using coarse-grained diffusive and anomalous diffusion models. (I) Simulated morphologies at 21 AEL with the diffusive (left panel) or anomalous diffusion (right panel) model, with parameters extracted from *in vivo* data. Simulations were initialized from an *in vivo* MT-inhibited Class IV neuron. The diffusive model fails to reproduce the observed morphology whereas the anomalous diffusion model accurately captures it (Compare left panel in I with right panel in A). (J,K) Morphometric parameters measured *in vivo* (black) and *in silico* (anomalous diffusion model in red, diffusive model in blue). In vivo, n = 4 neurons; *In silico*, n = 40 simulations. The anomalous diffusion model accurately reproduces both total arbor length (J) and semi-minor (K) observed *in vivo*, whereas the diffusive model does not. C-F,K,J,K. Data show mean ± SD.

To determine the impact of microtubule-mediated stabilization on long-term branch dynamics and its role in determining final neuronal shape, we simulated neuronal growth using parameters extracted from microtubule-depolymerized Class IV neurons (Movie S18). Simulations using the anomalous diffusion model accurately reproduced both the morphology and dynamical features of microtubule-inhibited Class IV neurons *in vivo* (Figure 6I-K). By contrast, simulations using the coarse-grained diffusive model, which does not incorporate α, largely overestimated both the dynamics and final morphology of microtubule-inhibited Class IV neurons (Figure 6J,K) These results show that dendritic branch dynamics are intrinsically subdiffusive, and that microtubule-mediated stabilization tunes the degree of subdiffusivity captured by the anomalous diffusion exponent α.

Microtubule-depolymerized Class IV neurons resemble Class I neurons as their size is strongly reduced and their dynamics is subdiffusive, with an anomalous diffusion exponent α close to 0.6. This suggests that microtubules are differently regulated and/or organized in these two neuronal classes. We therefore measured the rate of microtubule network expansion per branch tip (Section 9 of Supplementary Materials and Methods) and found that it was ∼5-fold higher in Class IV than in Class I neurons (0.027 µm.min-1 vs. 0.005 µm.min-1), indicating distinct microtubule dynamics (Table 15). Altogether, these results suggest that the divergent morphologies of Class I and Class IV neurons largely arise from differences in the local regulation of microtubules growth, which in turn control branch diffusivity over long time scales.

## Discussion

By exploring different classes of models encoding dendritic tree morphogenesis with measurements of the parameters in vivo, we mapped the morphospace of dendritic neurons. Systematic comparison between simulations and observations uncovered a minimal model defined by only four parameters of stochastic dynamics (λ, κ, σ, and α) that account for both neuronal morphogenesis and final shape. A single parameter, α, is necessary and sufficient to distinguish neuronal classes. Our work reveals that dendritic architecture emerges from the interplay of distinct cytoskeletal dynamics: actin-driven short-term fluctuations (σ), which drive dendritic arbor expansion and microtubule-dependent long-term dynamics (α), which set class-specific morphology.

Theoretical analysis of neuron front propagation shows that front speed is most sensitive to α over the relevant parameter range. Notably, the same parameter that distinguishes neuronal classes morphologies, also emerges as the dominant control parameter of global arbor expansion. More broadly, the model predicts a clear separation between short-term fluctuations (σ), and long-term anomalous diffusion behavior (α). This separation maps closely onto the respective contributions of actin and microtubules to branch dynamics, with actin driving exploratory branch tip fluctuations and microtubules controlling long-term stabilization and growth persistence.

Despite this agreement, the model does not capture the biphasic growth of Class I neurons between 16 and 21 h AEL. In its current formulation, α is estimated as an effective anomalous diffusion exponent over 1 hour time lags and is therefore treated as constant over this developmental time-window. Allowing α to vary in time could account for transition between successive phases of arbor expansion, but would require substantially larger datasets, as reliable estimation of α requires branch trajectories spanning at least one hour.

The dynamics of dendritic branch growth and retraction — characterized by alternating phases of extension and collapse — bear striking similarities with the dynamics of cell protrusion during adhesion-based motility, where cycles of actin polymerization and arrest drive membrane advance. This analogy is further substantiated by the dependence of dendritic branch growth on adhesion to collagen in the extracellular matrix: in collagenase-treated embryos, branches fail to extend despite ongoing actin-dependent dynamics (Movie S19, S20).

However, key differences distinguish dendritic branching from cell motility. During cell motility, the nucleus translocates with the advancing membrane front, leaving the total membrane area roughly constant. During dendritic tree growth, by contrast, the cell body remains stationary and the overall membrane area increases substantially, presumably through mobilization of internal membrane stores. Because branches are long, thin membrane tubules (on the order of a few hundred nanometres), diffusion is likely limiting for their growth, making active transport critical for their organization and dynamics. Consistent with this, vesicular transport from the ER-Golgi secretory pathway is required for dendritic growth and branching, as it provides the membrane and proteins necessary for branch extension, and when severely disrupted leads to impaired dendrite arborization^45–47^.

These considerations shed light on two key properties of the stochastic branching dynamics we uncovered. First, branch extension is sub-diffusive on long time-scales both in *wild-type* Class I and microtubule-depolymerized Class IV neurons. This sub-diffusivity may stem from rising membrane tension that limits extension: the Brownian ratchet mechanism of membrane protrusion relies on fluctuations in the plasma membrane and actin filaments^48–50^ and increased membrane tension reduces protrusion efficiency^51–53^. In this framework, the role of microtubules in restoring diffusive dynamics in Class IV neurons could reflect microtubule-dependent transport of membrane vesicles that fuels plasma membrane growth, thereby maintaining low tension and sustaining persistent branch extensions. Another possibility is that microtubules transport regulators of actin nucleation to branch tips, increasing the persistence of growth, thus promoting net branch extension. In light of this, the morphological differences between Class I and Class IV could stem from differential regulation of microtubules polymerization and/or differential transport of actin regulators by microtubules.

Second, the distinct morphologies of Class I and Class IV neurons arise principally from differences in the anomalous diffusion exponent α, which depends on microtubule regulation. This suggests that neuronal shape is largely an emergent, self-organized property, genetically encoded by distinct transcription factors and implemented through differential regulation of microtubule-based transport. More broadly, our findings support the idea that cytoskeleton-regulated stochastic programs constitute a general strategy for generating robust neuronal architectures, as also observed during axonal targeting in *Drosophila*^54^. Consistent with this view, two key transcription factors Abrupt and Knot have been shown to control Class I and Class IV dendrite morphogenesis through cytoskeleton regulation. Abrupt, which is specifically expressed in Class I neurons, limits their branching by influencing microtubule nucleation organization through regulation of Centrosomin^55^. Knot, expressed in Class IV neurons, regulates Spastin, a microtubule-severing protein promoting dendritic branch extension by generating new microtubules ends that act as additional nucleation sites for polymerization^42^. This points to different regulations of microtubules in Class I and Class IV neurons. It will be interesting in future work to characterize how microtubule regulation underlies differential diffusive vs subdiffusive dynamics in Class I and Class IV neurons. External cues may also contribute to specific features of dendritic arborization. In particular, the dorsal branch of Class I neurons grows in an invariant pattern, implying a still-unidentified directional cue.

Our approach is data-driven and comparative, exploring different models to identify the one that best encodes dendritic trees. The model we arrived at is parsimonious — four parameters — and highlights the singular role of the anomalous scaling exponent α and the underlying contribution of microtubules. In the framework proposed by David Marr^56^, our model operates at the algorithmic level of understanding: it reveals the low-dimensional logic by which a biological system accomplishes a task — here, the differential shaping of Class IV and Class I dendritic arbors. Each parameter is directly connected to observables, rendering the model both generic and transparent. The roles of actin and microtubules in tuning σ, which captures short term branch dynamics, and α, which reflects long-term anomalous diffusion dynamics, respectively, pertain to Marr’s implementational level. Our work paves the way for a more focused investigation of microtubules organization, polarity, and transport properties in regulating branch stochastic dynamics, long-term diffusivity, and their differential control between neuronal classes. More broadly, we advocate the use of such data-driven, phenomenological approaches and computational modelling to uncover how complex morphologies emerge from high-dimensional molecular processes.

## Supporting information

Summary tables

## Material and Methods

### *Drosophila* strains and genetics

Two distinct Gal4 lines were used to image Class-I neurons development: *rluv3-Gal4* (^55^ for primary branch elongation and *221-Gal4*^36^ for later stages. Both lines were recombined with *UAS-CD4::mNeonGreen*^14^. Imaging of Class-IV neurons was performed using *ppk-CD4::TdGFP*^57^. Visualization and quantification of actin and microtubules were achieved by crossing *rluv3-Gal4* or *ppk-Gal4*^58^ to *UAS-mNeonGreen:: αtub-T2A-mScarlet-I::utABD-P2A-CAAX::HALO*_7_ (this study). All (O’Donnell & Bernstein, 1988)^59^ to prevent muscular contractions of the embryos during imaging. All fly constructs and genetics are listed in the following table.

| Genotype | Figure |
| --- | --- |
| *w; mhc<sup>1</sup>/CyOGFP;rluv3-Gal4,UAS-CD4::mNeonGreen* | Fig 1D, J-M; Fig 2E-F; Fig 3D-F; Fig 5C; Fig S1A-F; Fig S2A-C, S2H-L; Fig S3A-L; Fig S4A-D; Fig S5D-F, J-L and Fig S10A, A' |
| *w; mhc<sup>1</sup>/CyOGFP;221-Gal4,UAS-CD4::mNeonGreen* | Fig 1D, E, J-M; Fig 2H, J-L; Fig 3F, J-L; Fig 4A-F; Fig S1A-F; Fig S4I-L, Q-T; Fig S6A, C, H-K; Fig S10B, B' |
| *w; mhc<sup>1</sup>/CyOGFP;ppk-CD4::TdGFP* | Fig 1B, C, F-I; Fig 2A-C, D, G-J, M-N; Fig 3A-C, F-I; Fig 4A-C, G-I; Fig 5D; Fig 6A-K; Fig S1G-L; Fig S2D-L; Fig S3M-X; Fig S4E-H, M-P, U-X; Fig S5A-C, G-I; Fig S6A, B, D-G |
| *w; mhc<sup>1</sup>/CyOGFP;rluv3-Gal4,UAS-mNeonGreen::αtub-T2A-mScarlet-I::utrABD-P2A-CAAX::HALO<sub>7</sub>* | Fig 5A-A''; Fig S8B, B''; Fig S9A-A'' |
| *w; mhc<sup>1</sup>,ppk-Gal4/CyOGFP; UAS-mNeonGreen::αtub-T2A-mScarlet-I::utrABD-P2A-CAAX::HALO<sub>7</sub>* | Fig 5B-B''; Fig S8C, C'; Fig S9B-B'' |

### Constructs and transgenesis

The pUASt-10x–mNeonGreen::αtub84B–T2A–mScarlet-I::utABD–P2A–HaloTag7x-CAAX Plasmid (Figure S8A), which was generated to simultaneously label microtubules, F-actin, and the plasma membrane in neurons, was assembled from three modular components. First, a membrane marker backbone containing seven tandem TagHalo7 repeats fused to a C-terminal CAAX motif was reconstructed in pBluescript II SK+ and modified by Gibson Assembly to introduce flanking BsmBI sites for subsequent Golden Gate cloning. Second, the GFP coding sequence of pJFRC81 10xUAS-IVS-Syn21-GFP-p10 (Addgene #36432) was replaced by In-Fusion cloning with a cassette encoding mNeonGreen–αTub84B and mScarlet-I–utABD with a T2A self-cleaving peptide ensuring bicistronic expression. Third, the full multi-reporter construct was assembled by quadripartite Golden Gate cloning (BsmBI) from a modified pJFRC81 backbone, two PCR-amplified fragments encoding the microtubule and actin reporters (BsmBI_Neon-αTub84B-GSG-T2A and BsmBI_mScarletI-utABD-GSG-P2A), and the HaloTag7x-CAAX donor plasmid. Internal BsmBI sites were removed by silent mutagenesis to ensure cloning compatibility, and GSG linkers were introduced upstream of the T2A and P2A peptides to improve downstream translation.

The final plasmid was verified by Sanger and whole-plasmid sequencing, and transgenic flies were generated by phiC31-mediated integration at attP2, attP40 and 24749 landing sites. FASTA sequences of all plasmids and detailed cloning strategy are available upon request.

### Sample preparation and embryos micro-injection

Flies were maintained in cages at 25 °C. Embryos were prepared as previously described^60^.In brief, embryos were collected on apple juice agar plates supplemented with yeast paste, dechorionated with 2.6% bleach for 90 s, transferred to a mesh basket and then rinsed thoroughly with distillated water before being transferred on a flat piece of clean agar. They were then staged (13 –16 h AEL) and selected under a dissection microscope and were aligned with their lateral side facing upward. For live imaging, Class I embryos were transferred to a glass coverslip coated with homemade glue and covered with Halocarbon 200 oil (Polyscience) to avoid drying during imaging. For long-term imaging of Class IV neurons, embryos were not glued due to the sensitivity of these neurons to mechanical perturbation. Instead, they were carefully transferred onto a coverslip using a brush and directly immersed in Halocarbon oil, with a piece of nylon mesh overlaid to stabilize them (Millipore, NY8H02500). For micro-injections of Class I and Class IV neurons, embryos were collected and aligned as described before, glued on a coverslip and systematically desiccated with Drierite for 9min30. Embryos were then covered with Halocarbon 200 oil and injected laterally with drugs or HaloTag ligand. Due to limited diffusion, injections were performed between the epidermal and muscles layers of the embryo, where neurons are developing.

### Drugs and Halo-tag ligand injections

For actin inhibition, a 100 mg/mL stock solution of Cytochalasin-D (Sigma-Aldrich, C8273) was prepared in DMSO, aliquoted, stored at −80 °C, and diluted 1:10 (final concentration: 10 mg/mL) in PBS prior to injection. For microtubules inhibition, a combination of two drugs was used. A 10 mg/mL stock solution of Vinblastine (Interchim, SSV610/11762) and a 25 mg/mL stock solution of Colcemid (Interchim, 11509B/15364) were prepared in DMSO, aliquoted, and stored at −80 °C. The injected solution consisted of a 1:12 dilution of Vinblastine (final concentration: 0.83 mg/mL) and a 1:6 dilution of Colcemid (final concentration: 4.16 mg/mL) in PBS. For membrane labeling, a 760 mM stock solution of HaloTag ligand JF646 (Promega, GA1120) was prepared in DMSO, aliquoted, stored at –80°C and diluted 1:5 in PBS prior to injection. All drugs and ligand were injected about 5 min before imaging. For ECM disruption, a 100mg/mL stock solution of collagenase (Merck C0130-100mg) for prepared in water, aliquoted and stored at –80C. The drug was injected without any dilution.

### Live imaging

Live imaging was performed at room temperature (21–22 °C) using either a spinning disc confocal (CSU-X1, Yokogawa) Nikon Eclipse Ti inverted microscope equipped with two cameras (Rolera EM-C^2^, Q-Imaging or Kinetix22, Photometrics; distributed by Roper) or with a Zeiss 880 confocal microscope equipped with GaAsP hybrid detectors.

Long-term time-lapse imaging of *rluv3>CD4::mNeonGreen* (early Class-I), *221>CD4:: mNeonGreen* (Class-I) and *ppk-CD4-Td::GFP* (Class-IV) was performed on the spinning disc confocal using a 491nm excitation laser and a 100X/1.45NA Plan Apo oil-immersion objective (Nikon). Z-stacks were acquired with a step size of 1μm for Class-I neurons and 0,5 μm for Class-IV neurons, spanning from the epidermis to the muscle layers, at a rate of one stack per minute for *221>CD4::mNeonGreen* and *ppk-CD4::TdGFP* and one stack every 5 min for *rluv3>CD4::mNeonGreen,* for at least 6h. Short-term time-lapse imaging was performed using similar settings with a frame rate of one stack every 5s for 10 min. Actin inhibition experiments were acquired with a frame rate of one stack per 30s for 10-15 min, microtubules inhibition and stabilization experiments were acquired with a frame rate of one stack per minute for 1-8h.

Live imaging of actin and microtubules in Class-I neurons (*rluv3> mNeonGreen::αtub, mScarlet-I::utABD, CAAX::HALO*_7_) was performed on the spinning disc confocal microscope with the 100x objective. Dual-color time-lapse imaging was achieved by simultaneous exciting with 491-nm (mNeonGreen::*α*Tub) and 561-nm (mScarlet-I::UtABD) lasers, using a dichroic mirror to collect emission signals on two cameras. Z-stacks of 1μm were acquired every minute for at least 6h. Imaging of Class-IV neurons (*ppk> mNeonGreen:: αtub, mScarlet-I::utABD, CAAX::HALO*_7_) was performed on the Zeiss 880 confocal microscope with a 63X/1.4 NA oil objective, using simultaneous excitation at 488-nm (mNeonGreen::*α*tub) and 561-nm (mScarlet-I::UtABD), with emission signals collected on GaAsp hybrid detectors. Z-stacks (1μm step size) were acquired every minute for at least 6h.

For imaging actin, microtubules and membrane following JF646 HaloTag ligand injection, triple-color images were acquired on the spinning disc confocal microscope using sequential excitation at 491-nm (mNeonGreen::*α*tub), 561-nm (mScarlet-I::UtABD) and 642-nm (CAAX::HALO_7_-JF646).

Imaging conditions (line averaging, camera exposure time, laser power) were optimized and kept constant across experiments. Image analysis and quantification were performed on maximum-intensity projections of z-stacks. Additional details on image analysis and computational modeling are provided in the Supplementary Material and Methods.

## Data availability

Summary statistics supporting the analyses are provided in the Supplementary Tables. The underlying processed datasets, including time-series measurements and branch-trajectory data, are available at this repository: https://github.com/MarcEricP/EncodingNeuronalShape. The raw imaging data are available from the corresponding author upon request.

## Code availability

Custom code for stastical analysis and simulations is available at this repository: https://github.com/MarcEricP/EncodingNeuronalShape. Custom code used for image analysis and branch tracking is available to editors and reviewers upon request. The image-processing pipeline will be released in a public repository linked to a methods preprint.

## Supplementary information

**Supplementary Material and Methods (see PDF document)**

**Supplementary information on front propagation theory (see PDF document)**

## Supplementary Figure legends

**<u>Figure S1</u>. Growth morphometrics of Class I and Class IV neurons**

(**A-L**) Growth morphometrics of Class I (green, **A-F**) and Class IV (magenta, **G-L**) neurons. (**A,G**) Total length. (**B,H**) Semi-minor axis (lateral extension). (**C,I**) Semi-major axis (vertical extension). (**D,J**) Aspect ratio. (**E,K**) Number of branching points. (**F,L**) Density. Solid lines indicate the mean; shaded area indicates mean ± 1 SD. n=5 Class IV neurons; n=6 (13h-16h) and n=7 (16h-21h) Class I neurons.

**<u>Figure S2</u>: Two-state Markov model quantifications**

(**A-F**) Trajectories selected for analysis from automated tracking over high-temporal resolution movies (1 frame every 5s for 10 min) at 16, 18 and 20h AEL. Class-I (**A-C**): 21 acquisitions, 332 trajectories, 1393 phases. Class-IV (**D-F**): 16 acquisitions, 780 trajectories, 3448 phases.

(**G**) Example of piecewise linear fitting. Models with increasing number of segments are iteratively tested and the optimal model is selected by minimizing the Bayesian Information Criterion (BIC), which balance fit quality and model complexity.

(**H-L**) Comparison of growth and shrinkage dynamics between Class I (green) and Class IV (pink) neurons. (**H,I**) Comparison of v_on_ (**H**) and v_off_ (**I).** Asterisks denote significance based on a Welch t-test with Holm correction: (**H**) 16h p = 0.84; 18h p = 0.19; 20h p = 5.8e-11; (**I**) 16h p = 0.27; 18h p = 9.2e-12; 20h p = 2.8e-35 (**J**) Example of merged phases of identical type (growth in orange, shrinkage in blue) obtained from the optimal piecewise linear fit (**K,L**) Comparison of τ_on_ (**K**) and τ_off_ (**L**). Welch t-test with Holm correction: (**K**) 16h p = 0.67; 18h p = 0.27; 20h p = 0.73; (**L**) 16h p = 0.10; 18h p = 2.87e-8; 20h p = 1.2e-5.

**<u>Figure S3</u>. Distributions of phases velocity and duration**

(**A-L**) Class-I neurons. (**M-X**) Class-IV neurons. (**A-D;M-P**) 16h AEL. (**E-H;Q-T**) 18h AEL. (**I-L;U-X**) 20h AEL. (**A,E,I,M,Q,U**) v_on_, (**B,F,J,N,R,V**) v_off_, (**C,G,K,O,S,W**) τ_on_,(**D,H,L,P,T,X**) τ_off_.

**<u>Figure S4</u>. Comparison of morphometrics in data versus simulations**

(**A-H**) Two-state Markov model: (**A-D**) Class-I; (**E-H**) Class-IV.

(**I-P**) Coarse-grained diffusive model: (**I-L**) Class-I.;(**M-P**) Class-IV.

(**Q-X**) Anomalous diffusion model: (**Q-T**) Class-I; (**U-X**) Class-IV.

(**A-X**) *In vivo*: n = 7 (Class I); n = 5 (Class IV). *In silico:* n = 70 (Class I) and n = 50 (Class IV). Data show mean ± SD.

**<u>Figure S5</u>. Phase velocity and duration are not independent**

(**A-F**) Joint distributions of phase velocity and phase duration at 16 h AEL (**A,D**), 18 h AEL (**B,E**) and 20 h AEL (**C,F**). Each dot represents a single phase (growth in blue, shrinkage in orange). Solid lines show linear regressions of velocity versus log(duration), revealing a coupling between the two variables. (**A-C**) Class-IV neurons: growth, 16h: p = 1.0e-6; 18 h: p = 2.8e-60; 20h: 1.6e-50. Shrinkage, 16h: p =7.5e-4; 18 h: p = 5.0e-64; 20h: 5.4e-50. (**D-F**). Class-I neurons: growth, 16h: p = 9.3e-6; 18 h: p = 5.1e-14; 20h: 2.6e-12. Shrinkage, 16h: p = 2.1e-8; 18 h: p = 8.8e-23; 20h: 3.7e-18.

(**G-L**) Comparisons of mean displacement of branch-tip trajectories from experimental data (black), simulations from the 2-state Markov model assuming independence between velocity and duration (purple), and a model incorporating velocity-duration coupling (cyan), in Class IV (**G-I**) and Class I (**J-L**) neurons. Diffusion coefficients D are reported with bootstrap 95% confidence intervals. P values are obtained from two-sided bootstrap tests on ΔD (B = 200 resamples).

**<u>Figure S6</u>. Coarse-grained diffusive model with drift**

**(A)** Evolution of drift with time, measured as the mean of the distribution of increments. Solid lines indicate the mean; shaded area indicates mean ± 1 SD. n=5 Class IV neurons; n=7 Class I neurons. (**B,C**) Comparison of Class IV (**B**) and Class I (**C**) neuronal morphologies at 21h AEL (left panels) with simulated neurons (right panels). (**D-K**) Morphometric parameters measured in *vivo* (black) and *in silico* using the coarse-grained diffusive model (blue) or the coarse-grained diffusive model with drift (cyan), for Class IV (**D-G**) and Class I (**H-K**) neurons. Including drift maintains accurate predictions for Class IV neurons but fails to recapitulate Class I morphology. (**D-K**) *In vivo*: n = 7 (Class I); n = 5 (Class IV). *In silico:* n = 70 (Class I) and n = 50 (Class IV). Data show mean ± SD.

**<u>Figure S7</u>. Variation of anomalous diffusion model parameters *in silico***

(**A-P**). Morphometrics measurements from simulations in which a single parameter of the anomalous diffusion model is varied with all other parameters are fixed to their class-specific measured values. Data show mean and SD, each dot represents one simulation realization. Panels: (**A-D**) Total length; (**E-H**) Semi-minor axis; (**I-L**) Number of branching points; (**M-P**) Density. Parameters variations: (**A,E,I,M**) λ varies; (**D,F,J,N**) κ varies; (**C,G,K,O**); σ varies; (**D,H,L,P**) α varies. For λ, σ and κ, the x-axis represents the multiplicative factor relative to the reference value (x=1).

**<u>Figure S8</u>. A new tool to visualize actin, microtubules and the plasma membrane**

**(A)** Schematic of the plasmid pUASt-10x–mNeonGreen::αtub84B–T2A–mScarlet-I::utABD–P2A–HaloTag7x-CAAX built to simultaneously visualize actin, microtubules and the plasma membrane. **(B,C)** Live images of Class I (**B,B’**) and Class IV (**C,C’**) neurons expressing the construct. *Rluv-gal4* and *ppk-gal4* were used to drive expression of the construct in Class I and Class IV neurons, respectively. Microtubules (MTs, mNeonGreen) are show in green, actin in magenta (mScarlet-I) and the plasma membrane in yellow (injection of JF646 HALO-tag ligand). In both neuronal types, actin is enriched at dynamic dendritic tips whereas microtubules are enriched in proximal dendritic regions.

**<u>Figure S9</u>. Evolution of actin and microtubule networks over time**

(**A,B**) Zoom-in views of timelapse-imaging of individual branches in Class I (**A-A”**) and Class IV (**B,B”**) neurons. Actin and microtubule networks are in magenta and green, respectively.

**Figure S10. Microtubules inhibition in Class I neurons**

(**A,A’**) Time-lapse imaging of Class I primary branch growth in control (**A**) or following injection of Colcemid and Vinblastine (**A’**). Microtubules inhibition impairs primary branch elongation.

(**B, B’**) Time-lapse imaging of Class I secondary branches growth in control (**B**) or following injection of Colcemid and Vinblastine (**B’**). Microtubules inhibition prevents stabilization of secondary branches.

## List of supplementaryl movies

**Movie S1**: Comparison of Class I and Class IV morphogenesis

**Movie S2**: Growth of Class I primary branch

**Movie S3**: High-temporal-resolution tracking of Class I branches

**Movie S4**: High-temporal-resolution tracking of Class IV branches

**Movie S5**: Simulation of Class I growth with the two-state Markov model

**Movie S6**: Simulation of Class IV growth with the two-state Markov model

**Movie S7**: Simulation of Class IV growth with the coarse-grained diffusive model

**Movie S8**: Simulation of Class I growth with the coarse-grained diffusive model

**Movie S9**: Simulation of Class I growth with the anomalous diffusion model

**Movie S10**: Simulation of Class IV growth with the anomalous diffusion model

**Movie S11**: Evolution of actin and tubulin networks overtime in Class I neurons

**Movie S12**: Evolution of actin and tubulin networks overtime in Class IV neurons

**Movie S13:** Actin inhibition in Class IV neurons

**Movie S14:** Actin inhibition in Class I neurons

**Movie S15:** Microtubules inhibition in Class IV

**Movie S16:** Microtubules inhibition during primary branch elongation of Class I neurons

**Movie S17:** Microtubules inhibition during secondary branch elongation of Class I neurons

**Movie S18:** Simulation of microtubule-inhibited Class IV growth with the anomalous diffusion model

**Movie S19:** Effects of collagenase injection in Class I neurons

**Movie S20:** Effects of collagenase injection in Class IV neurons

## List of supplementaryl tables

Table 1: Class-IV dendritic morphometric measurements over time.

Table 2: Class-I dendritic morphometric measurements over time.

Table 3: Branch trajectory dataset used to quantify local growth and shrinkage dynamics.

Table 4: Branch-tip growth and shrinkage parameters in WT Class I and Class IV neurons.

Table 5: Comparison of branch-tip growth and shrinkage parameters between WT Class I and Class IV neurons.

Table 6: Temporal changes in branching rate λ across WT Class I, WT Class IV, and MT-inhibited Class IV neurons.

Table 7: Pairwise comparisons of branching rate λ between WT Class I, WT Class IV, and MT-inhibited Class IV neurons.

Table 8: Post-contact branch retraction amplitude κ in WT Class I and Class IV neurons.

Table 9: Comparison of post-contact branch retraction amplitude κ between WT Class I and Class IV neurons.

Table 10: Relationship between branch-tip velocity and phase duration.

Table 11: Diffusion coefficients estimated from experimental and simulated branch trajectories.

Table 12: Temporal evolution of short-term branch-length fluctuations σ _1min_ in WT Class I, Class IV neurons, and MT-inhibited Class IV neurons.

Table 13: Comparison of short-term branch-length fluctuations σ _1min_ between WT Class I, Class IV, and MT-inhibited Class IV neurons.

Table 14: Anomalous diffusion exponent α estimated from long-term branch trajectories in WT Class I and Class IV neurons.

Table 15: Average microtubule network expansion rate per branch tip in Class I and Class IV neurons.

## Acknowledgments

We thank all members of the Lecuit lab for stimulating discussions and feedback during the course of this project. We thank the imaging facility at IBDM, member of the National Infrastructure France-BioImaging (https://ror.org/01y7vt929) supported by the French National Research Agency (ANR-24-INBS-0005 FBI BIOGEN), for assistance with maintenance of the microscopes; FlyBase for maintaining curated databases; Daniel Cox, Yuh Nung Jan, Adrian Moore and Bloomington Drosophila Stock Center for providing y stocks. This work was supported by the ANR Dendromorph (AAP2022) to TL and JFR and received funding from France 2030, the French Government program managed by the French National Research Agency (ANR-16-CONV-0001) and from Excellence Initiative of Aix-Marseille University – A*MIDEX “through CENTURI. MEP was supported by a PhD fellowship of CENTURI and an ATER fellowship from the Collège de France. MC was supported by a long-term fellowship from EMBO (ALTF 62-2020), followed by – CENTURI and the ANR Dendromorph. CB and JFR are supported by CNRS and TL by the Collège de France.

## Author contributions

TL conceived the project. TL, CB and JFR co-supervised the study. CB and MC performed the live imaging experiments of Class I and Class IV neurons respectively. CB, EDS and MC performed drug injections experiments. MEP developed all the image processing algorithms, performed image analysis and quantifications, developed the simulations and conducted the theoretical work. EDS and JMP designed and generated all molecular constructs. All authors discussed the data. CB, MEP and TL wrote the manuscript and all authors made comments.

