## Supplementary figures and images for "Encoding neuronal shape in the stochastic dynamics of branching processes"

### Supplementary figures

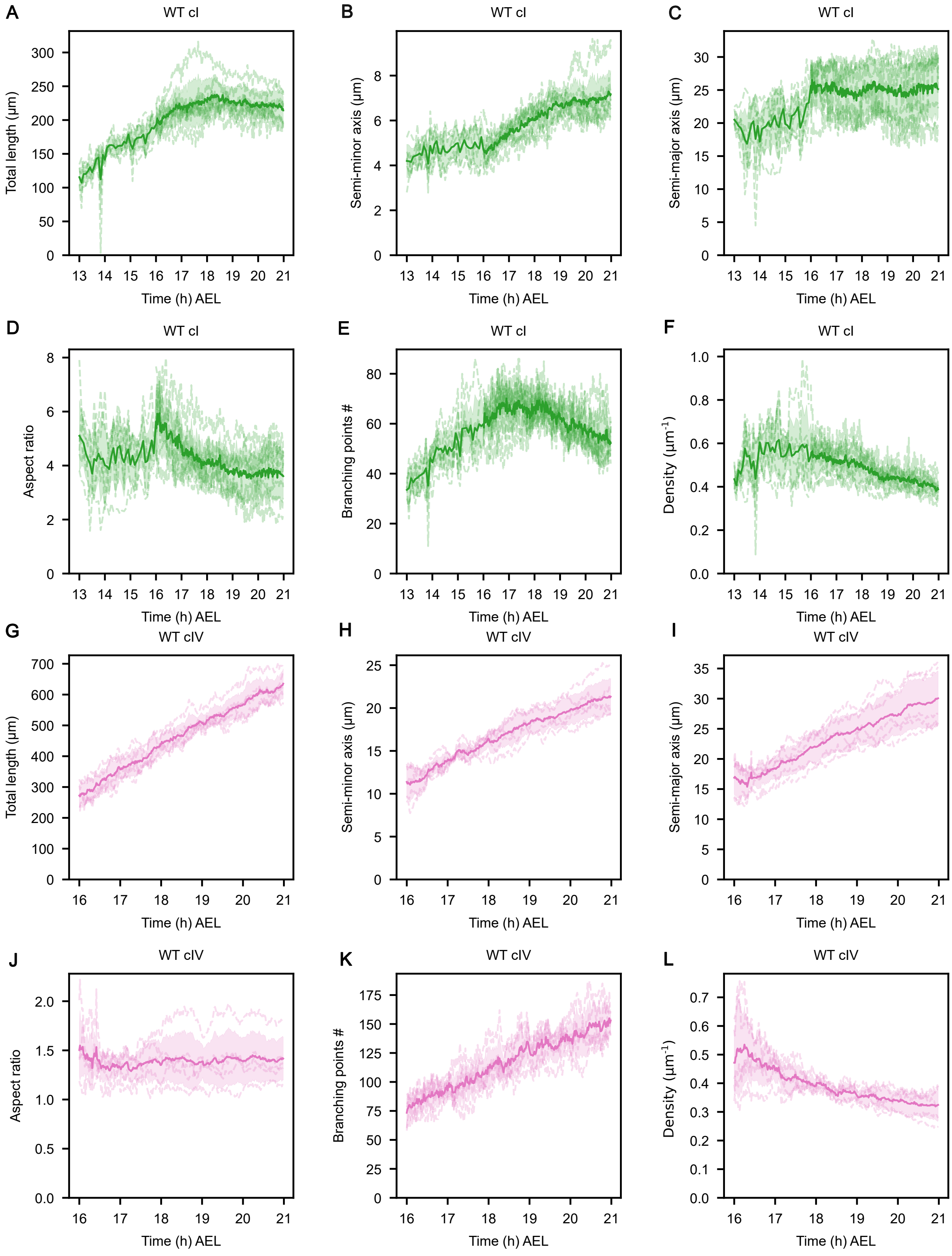

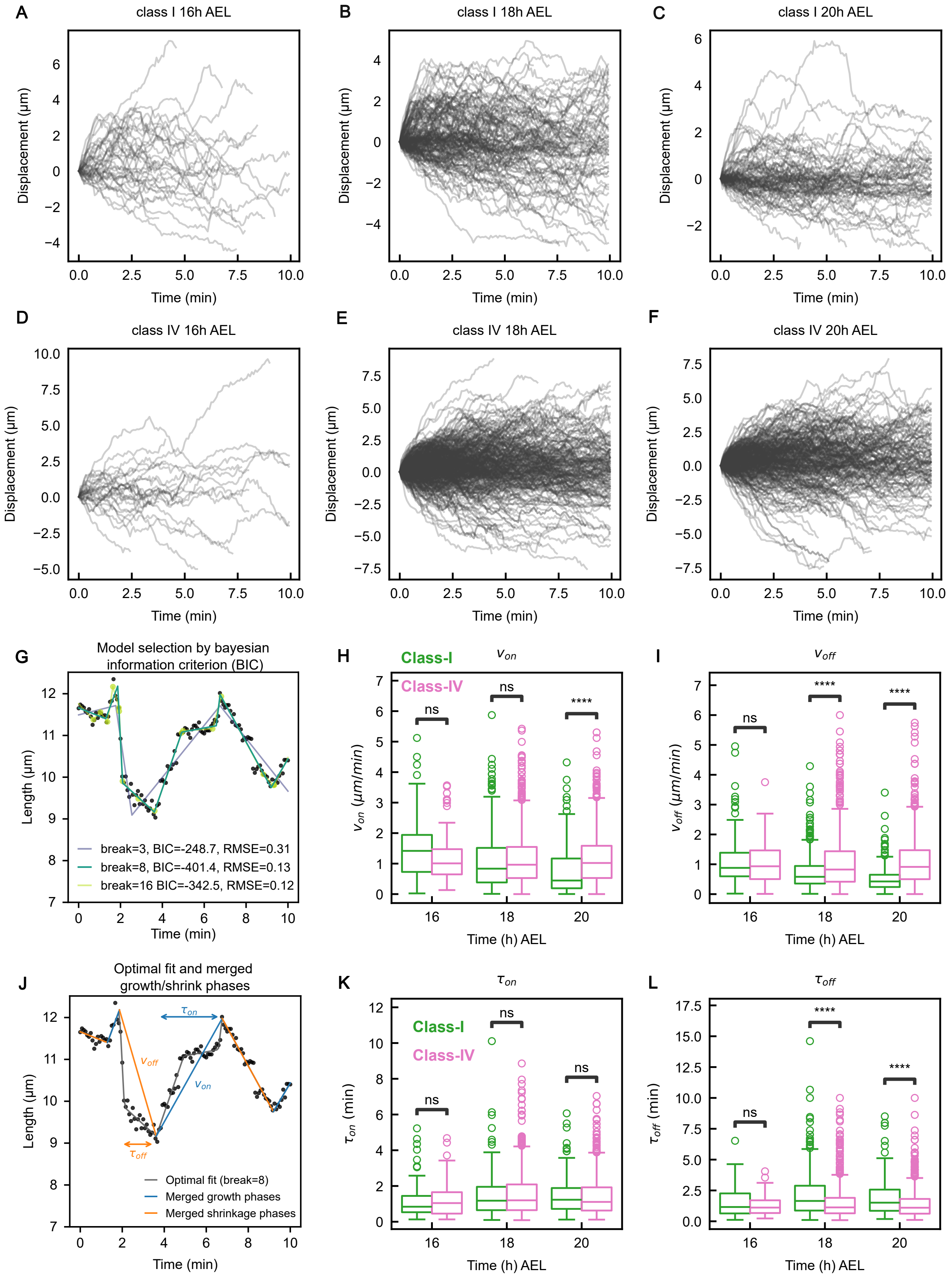

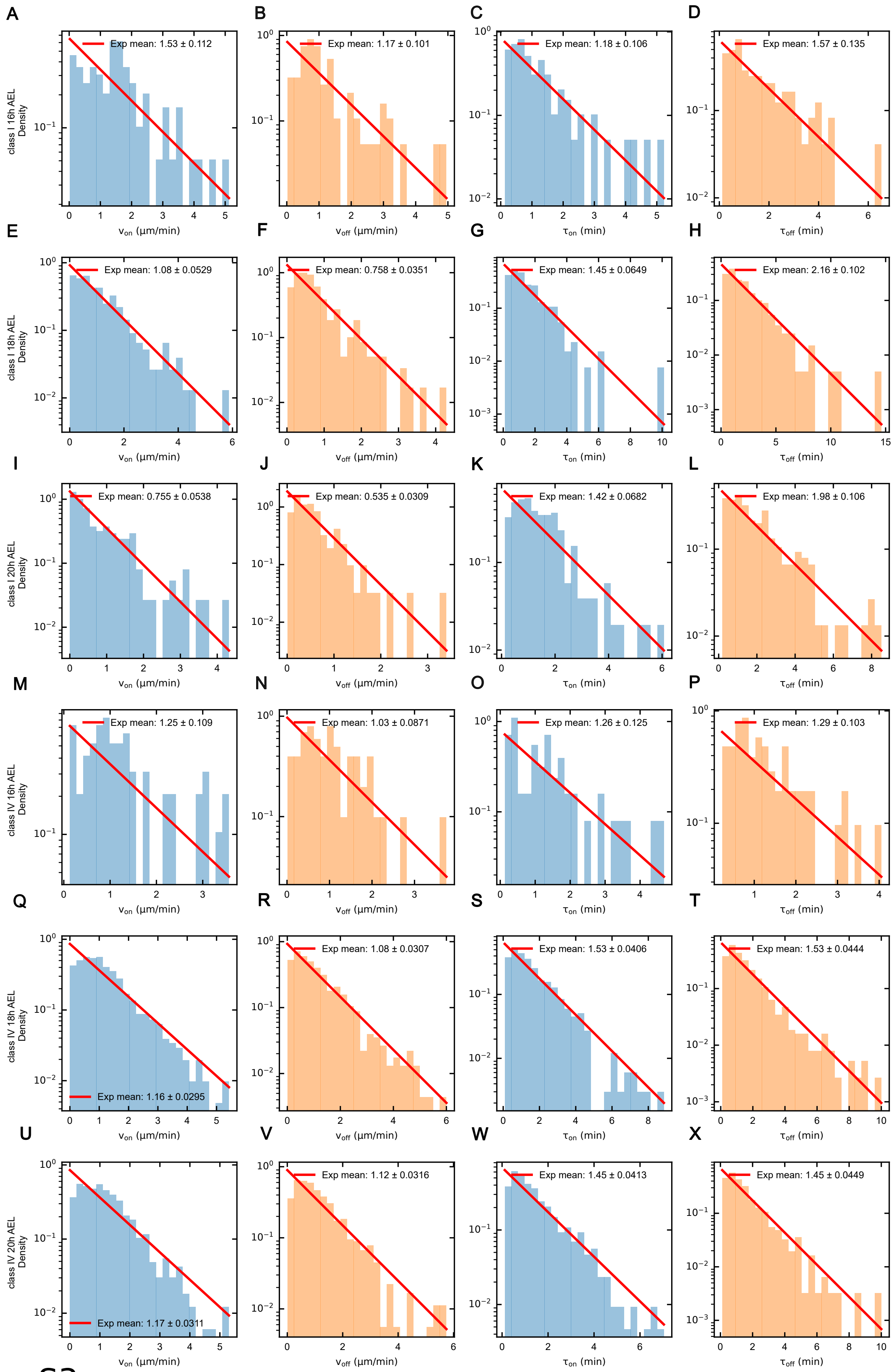

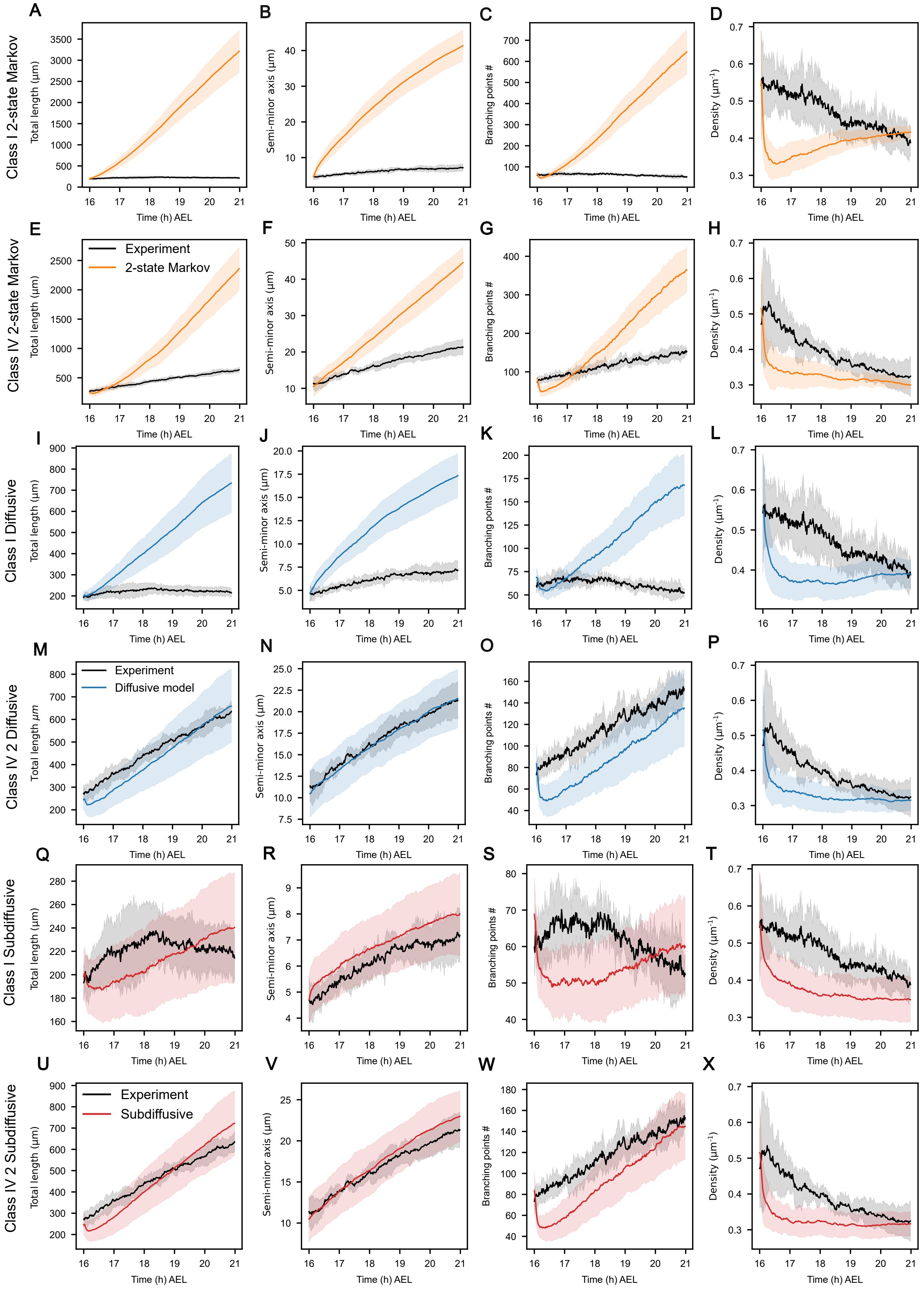

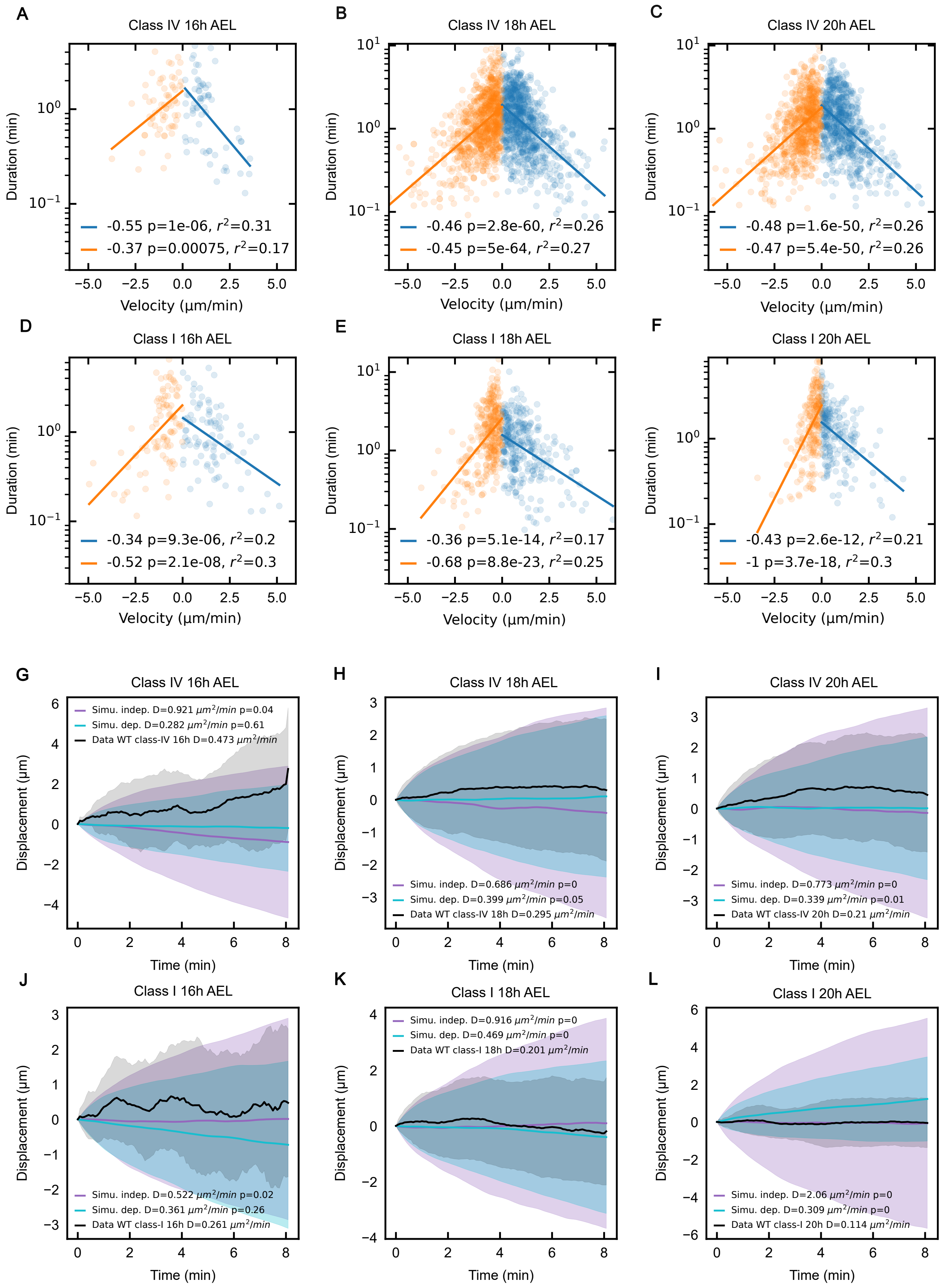

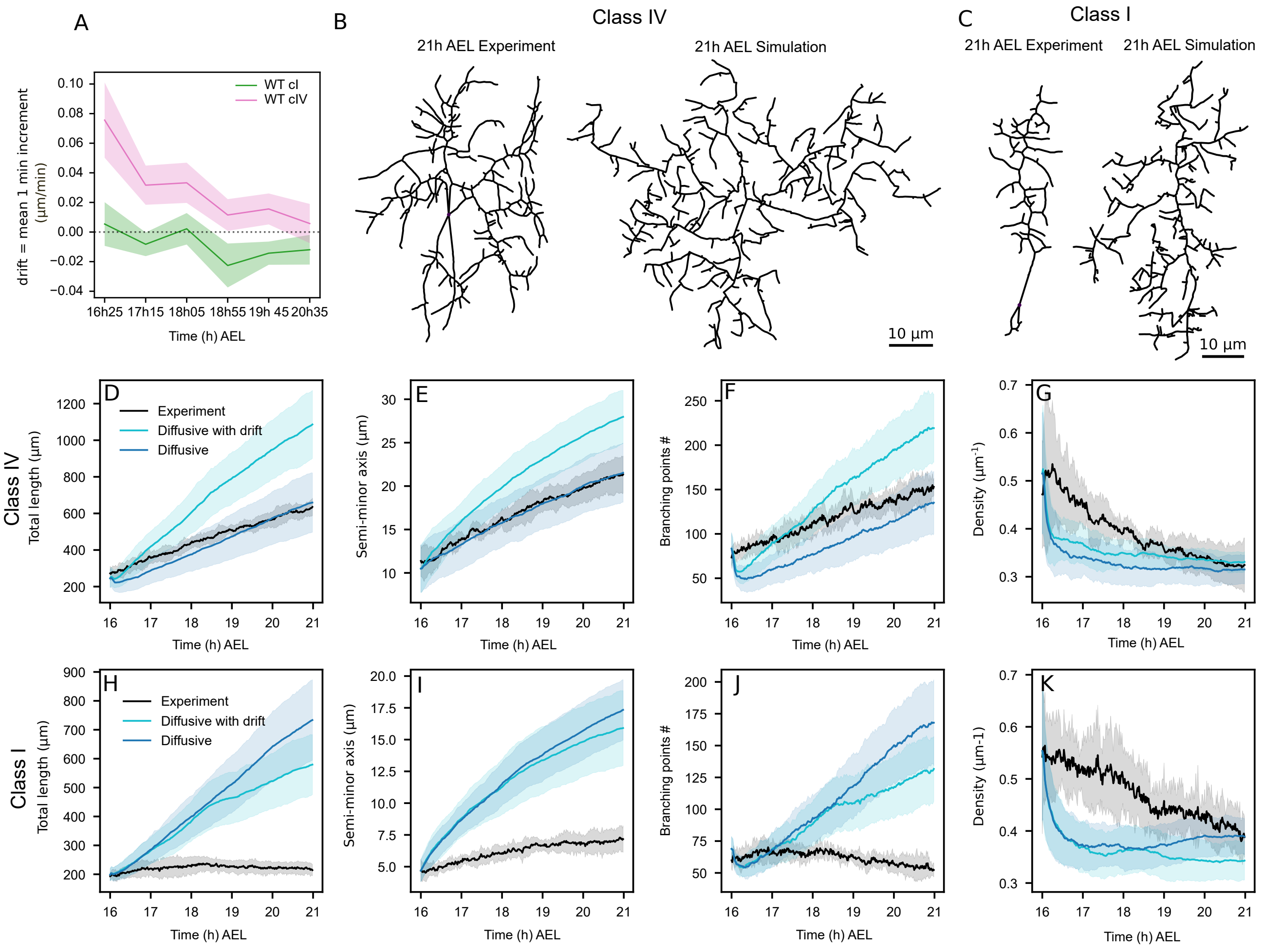

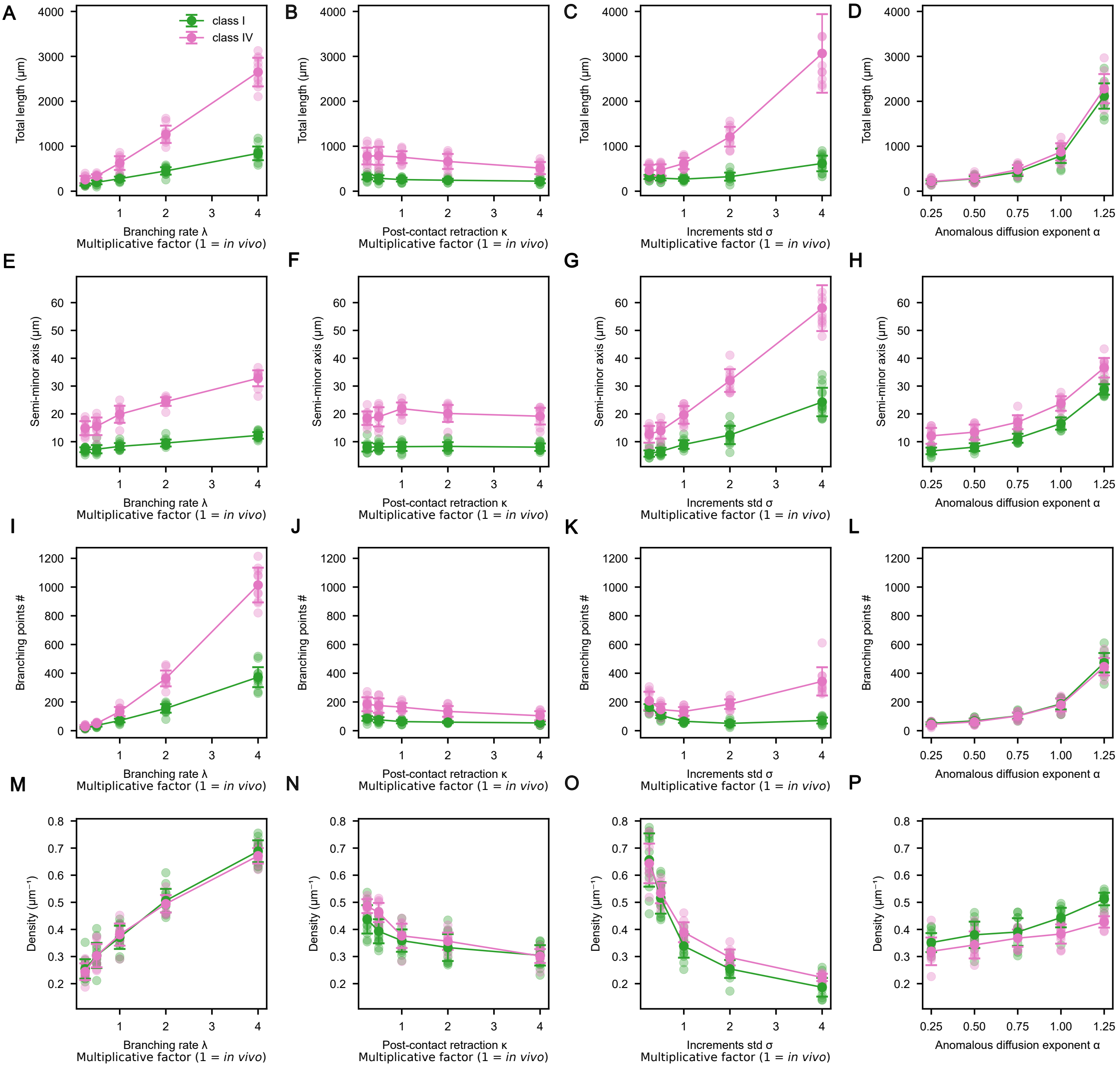

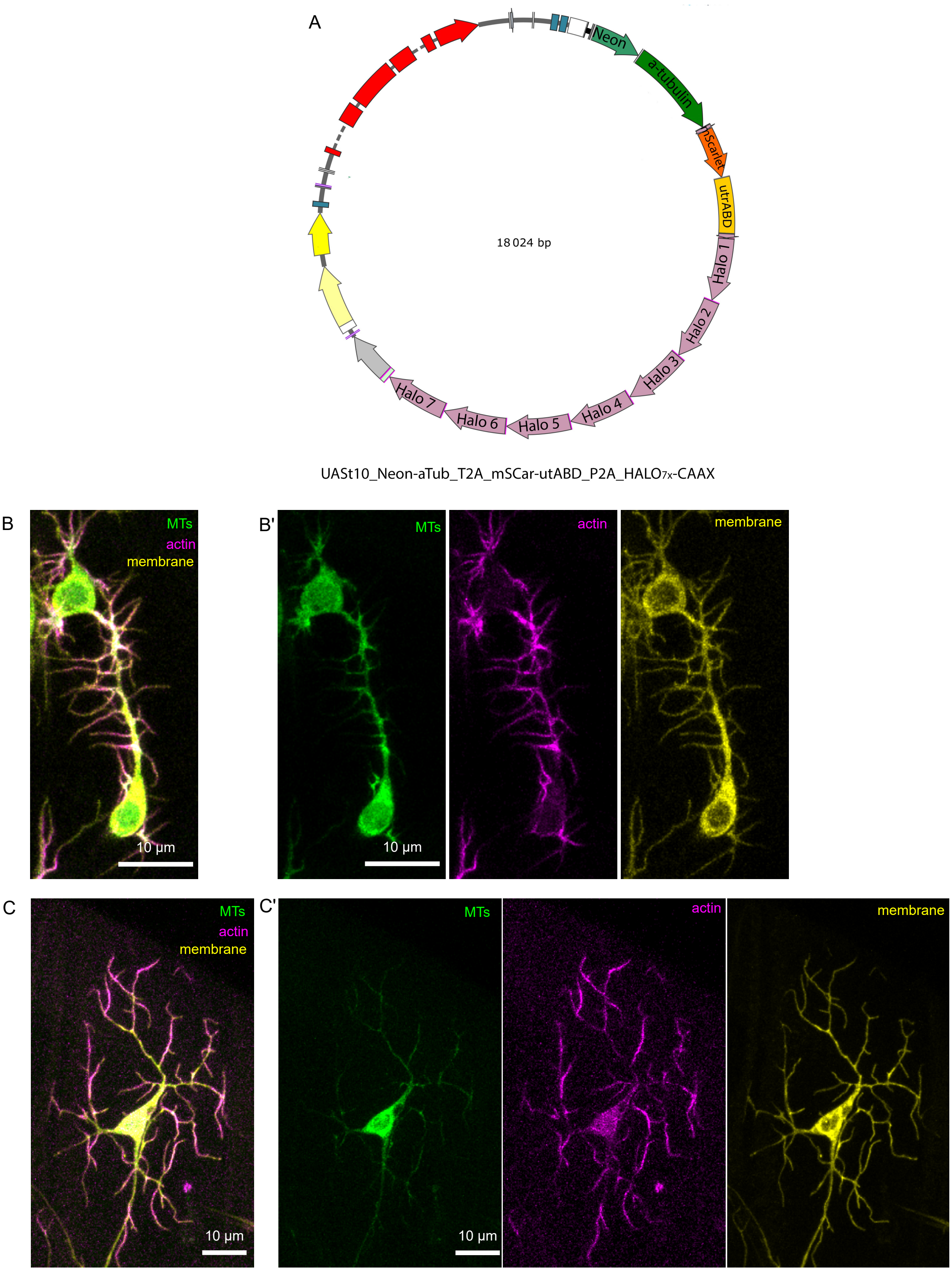

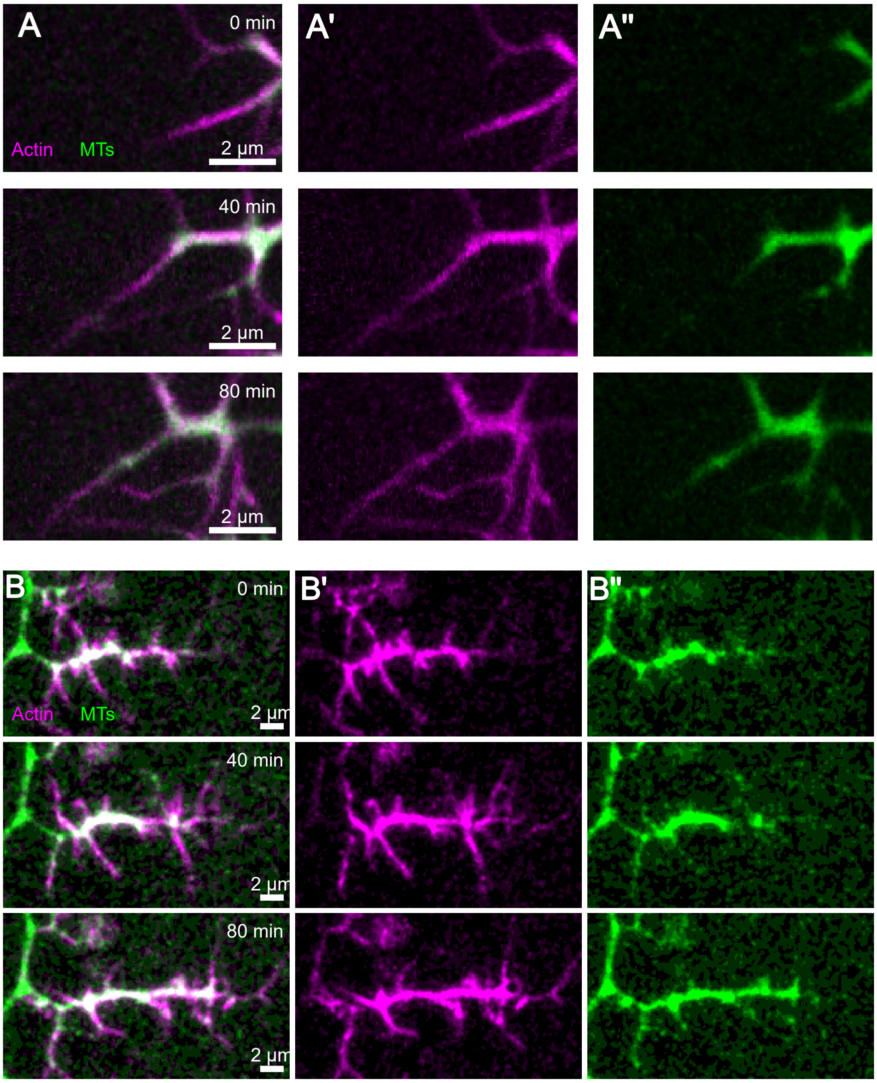

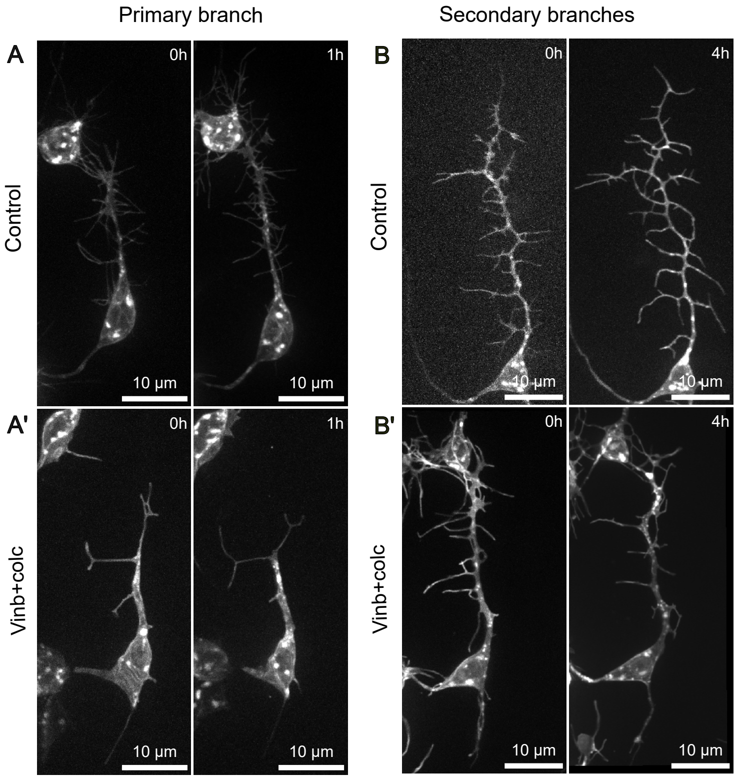
