## Supplementary information for "Encoding neuronal shape in the stochastic dynamics of branching processes"

May 5, 2026

[1] Aix Marseille Univ, CNRS, IBDM (UMR 7288), Turing Centre for Living Systems, Marseille, France.

[2] Aix Marseille Univ, CNRS, LAI (UMR 7333), Turing Centre for Living Systems, Marseille, France.

[3] Collège de France, 11 Place Marcelin Berthelot, Paris, France.

#### 1 Front speed and side branching

Previous work showed that tip-splitting branching random walks generate Fisher-KPP-type invasion fronts [1, 2]. In this model, new branch tips appear by tip splitting, and tips do not interact with each other after birth (Figure SI1A). With $D$ the diffusion coefficient and $r$ the splitting rate, with dimension $[r] = T^{-1}$, the density of tips $u$ follows

$$\partial_t u = D\partial_{xx} u + ru, \tag{1}$$

and the density front propagates with a constant minimal speed:

$$c^* = 2\sqrt{Dr}. \tag{2}$$

Hannezo et al. applied this model to various tip-splitting branching morphogenesis systems (e.g. mammary glands) [3]. However, neurons differ in two key respects: branching occurs via side branching along dendrites (Figure SI1B) rather than by tip splitting (Figure SI1A), and branch-tip motion can be subdiffusive rather than diffusive. Ouyang et al. considered a mean-field model of side branching with Markovian tip switching between growing, shrinking, and paused states [4]. For the full stochastic model, they derived an implicit front-speed selection equation, but not a simple closed-form expression or scaling law for the front speed as a function of the local dynamics parameters.

We replace tip splitting, represented by the term $ru$, with side branching. The key change is that new tips are no longer created locally at position $x$, but instead are generated from branches whose tips are located ahead of $x$. This is modeled by the lineic branching rate $\lambda$, with dimension $[\lambda] = L^{-1}T^{-1}$, and a nonlocal creation term $\int_x^\infty u$. To estimate how this mechanism changes the propagation velocity, we consider the equation

$$\partial_t u = D\partial_{xx} u + \lambda \int_x^\infty u. \tag{3}$$

Differentiating with respect to $x$ removes the nonlocal term and gives

$$\partial_{tx} u = D\partial_{xxx} u - \lambda u. \tag{4}$$

We then search for traveling-front solutions with exponential decay,

$$u(x, t) \sim e^{-\gamma(x-ct)}, \tag{5}$$

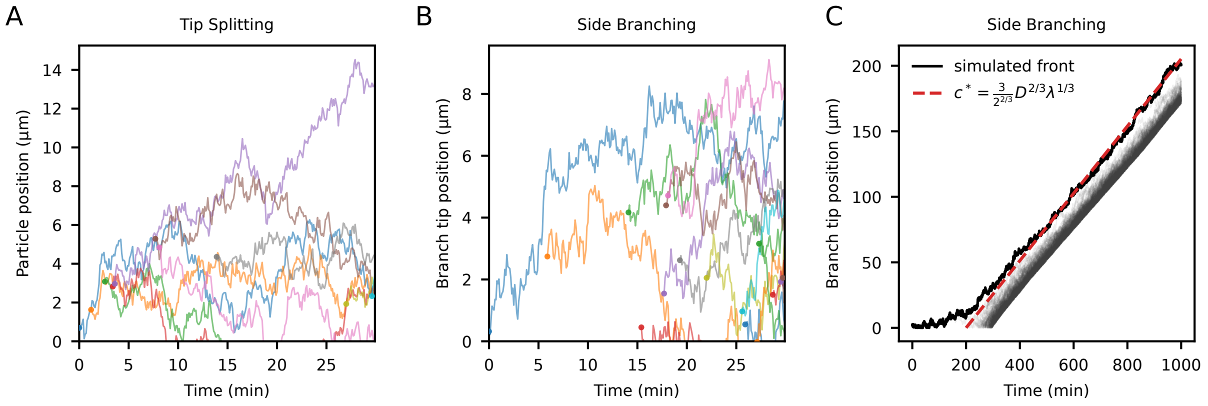


Figure SI1: (A) Example of a branching random walk with tip splitting: existing tips split at rate $r$ into two identical tips that have the same position and properties. Tips do not interact with each other. In this case, the minimal front speed is $c = 2\sqrt{Dr}$, where $D$ is the diffusion coefficient. (B) Branching random walk with side branching: new tips are born from the side of existing branches at lineic rate $\lambda$. (C) Theoretical front speed (dashed red line) against a 1D simulation (shaded curves indicate individual particles, solid curve indicates front position). Trajectories that are too far behind the front are not considered in order to accelerate simulations.

where $\gamma$ is the inverse decay length and $c$ is the front speed. Substituting this ansatz into the previous equation yields

$$-\gamma^2 cu = -\gamma^3 Du - \lambda u, \tag{6}$$

which directly gives the dispersion relation

$$c(\gamma) = \gamma D + \frac{\lambda}{\gamma^2}. \tag{7}$$

The selected velocity corresponds to the minimum of $c(\gamma)$ with respect to $\gamma$. Differentiating, we obtain

$$c'(\gamma) = D - 2\frac{\lambda}{\gamma^3}, \tag{8}$$

and the extremum condition therefore reads

$$\gamma_{|c'(\gamma)=0} = \left(\frac{2\lambda}{D}\right)^{1/3}. \tag{9}$$

Replacing this value into the dispersion relation gives the asymptotic front speed

$$c^* = \frac{3}{2^{2/3}} D^{2/3} \lambda^{1/3}. \tag{10}$$

This first result highlights the increased importance of tip motion in the side-branching case, where $c^*$ scales with $D^{2/3}$, whereas it scales with $D^{1/2}$ in the tip-splitting case (Figure SI1). Our results showed that the main difference in the local dynamics of class I and class IV neurons is that class I neurons exhibit subdiffusive behavior. We therefore introduce subdiffusion into the model in the next section.

### 2 Subdiffusive case

#### 2.1 Equation

Front speed in the context of anomalous diffusion was studied for Fisher-KPP equation [5, 6], but not with the non-local birth term induced by side-branching which we introduced in the previous section.

We derive the asymptotic front speed in the context of side branching, assuming that daughter branch tips do not inherit their parent history. Branch increments follow an $ARFIMA(0, \frac{\alpha-1}{2}, 0)$ process with $\Delta t$-increment standard deviation $\sigma$ and anomalous diffusion exponent $\alpha$ (see Materials and Methods) [7]. In this context, equation (3) can be adapted using a memory-kernel equation:

$$\partial_t u(x,t) = \int_0^t M(t-\tau)\partial_{xx} u(\tau, x)\, d\tau + \lambda \int_x^\infty u(t,y)\, dy. \tag{11}$$

For an $ARFIMA(0, \frac{\alpha-1}{2}, 0)$ process, the increment autocovariance decays as

$$E[X_t\, X_{t+n\Delta t}] \propto n^{\alpha-2}, \tag{12}$$

where $n$ is the number of steps, and the increments are drawn from a distribution with mean equal to zero. At large times, we assume that only the asymptotic power-law memory exponent matters. We therefore assume, as a phenomenological large-time approximation rather than a microscopic derivation, that the ARFIMA long-time memory can be represented by a fractional memory kernel with the same asymptotic power-law exponent:

$$M(t) \underset{t\to\infty}{\sim} \frac{K_\alpha}{\Gamma(\alpha-1)}\, t^{\alpha-2}, \tag{13}$$

where $K_\alpha \sim \frac{\sigma^2}{2\Delta t^\alpha}$ is the generalized diffusion coefficient.
Hence, for large times,

$$\int_0^t M(t-\tau)\, \partial_{xx} u(x,\tau)\, d\tau \sim \frac{K_\alpha}{\Gamma(\alpha-1)} \int_0^t (t-\tau)^{\alpha-2}\, \partial_{xx} u(x,\tau)\, d\tau = K_\alpha\, D_t^{1-\alpha} \partial_{xx} u(x,t), \tag{14}$$

with $D_t^{1-\alpha}$ the Riemann-Liouville derivative:

$$D_t^{1-\alpha} f(t) = \frac{1}{\Gamma(\alpha-1)} \int_0^t (t-\tau)^{\alpha-2} f(\tau)\, d\tau. \tag{15}$$

This results in the following large-time equation:

$$\partial_t u \sim K_\alpha D_t^{1-\alpha} \partial_{xx} u + \lambda \int_x^\infty u. \tag{16}$$

#### 2.2 Front speed

We compute the front speed using a similar approach to that in Section 1:
The ansatz

$$u(x,t) \sim e^{-\gamma(x-ct)} \tag{17}$$

gives

$$-\gamma^2 c u \sim -K_\alpha (\gamma c)^{1-\alpha} \gamma^3 u - \lambda u. \tag{18}$$

We define

$$F(c,\gamma) = c - K_\alpha (\gamma c)^{1-\alpha} \gamma - \frac{\lambda}{\gamma^2} \sim 0. \tag{19}$$

We are looking for $\gamma^*$ that minimizes $c(\gamma)$, and we denote $c^* = c(\gamma^*)$. At this point, $\frac{dc}{d\gamma}\Big|_{\gamma^*} = 0$, and since $F(c,\gamma) \sim 0$,

$$\frac{dF}{d\gamma} = \frac{\partial F}{\partial c}\frac{dc}{d\gamma} + \frac{\partial F}{\partial \gamma} \sim 0. \tag{20}$$

Therefore,

$$\left.\frac{\partial F}{\partial \gamma}\right|_{\gamma^*} = -(2-\alpha)K_\alpha(\gamma^* c^*)^{1-\alpha} + 2\frac{\lambda}{(\gamma^*)^3} \sim 0, \tag{21}$$

and

$$(\gamma^*)^{4-\alpha} \sim \frac{2\lambda}{(2-\alpha)K_\alpha c^{*(1-\alpha)}}. \tag{22}$$

Substituting into (19) yields

$$c^* \sim \Xi(\alpha) K_\alpha^{\frac{2}{2+\alpha}} \lambda^{\frac{2-\alpha}{2+\alpha}}, \tag{23}$$

with

$$\Xi(\alpha) = \frac{(4-\alpha)^{\frac{4-\alpha}{2+\alpha}}}{2^{\frac{2}{2+\alpha}}(2-\alpha)^{\frac{2-\alpha}{2+\alpha}}}. \tag{24}$$

With $K_\alpha = \frac{\sigma^2}{2\Delta t^\alpha}$,

$$c^* \sim \Xi(\alpha)\left(\frac{\sigma^2}{2\Delta t^\alpha}\right)^{\frac{2}{2+\alpha}} \lambda^{\frac{2-\alpha}{2+\alpha}}. \tag{25}$$

From the ansatz (Eq. 17), the characteristic time is $\frac{1}{\gamma^* c^*}$. The large-time approximation is therefore valid for $\Delta t \ll \frac{1}{\gamma^* c^*}$, which can be written as

$$\Delta t \sqrt{\frac{4-\alpha}{2-\alpha}\lambda c^*} \ll 1. \tag{26}$$

The front velocity is computed numerically for a range of parameters that satisfies this condition in Figure SI2C.

**1D Simulations** We compared the theoretical expression for the front speed with the front speed computed in one-dimensional simulations in which tips do not interact with each other after birth. As we were interested only in the front speed, we kept track of only the 5000 tips closest to the front, as tips farther away were very unlikely to have lineages that ever reach the front (Figure SI1C). We observed good agreement between simulations and theory over several orders of magnitude (Figure SI2A,B).

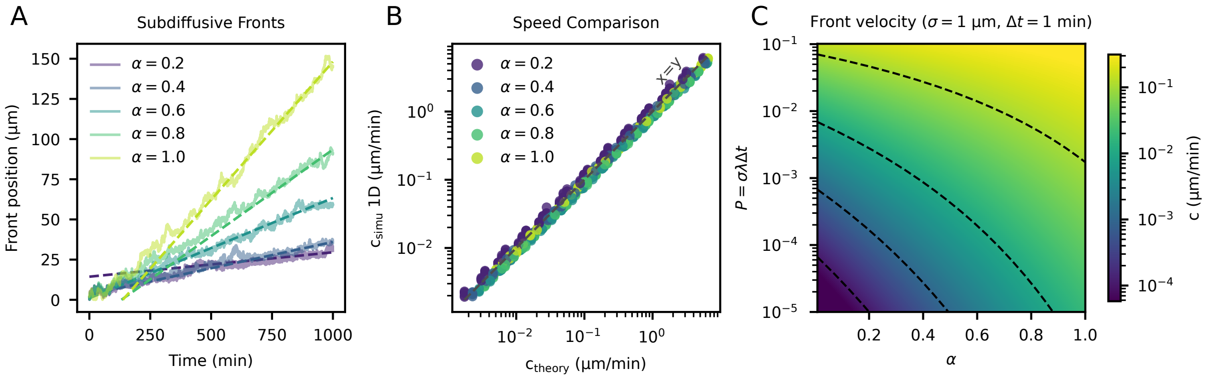


Figure SI2: (A) Comparison of 1D simulations and theory for the front with various values of $\alpha$. (B) Comparison of theoretical and simulated front speed for various values of $\alpha$, $\lambda$, and $\sigma$. For each value of $\alpha$, $\sigma$ ranges from $7.8125 \times 10^{-3}\,\mu\text{m}$ to $8\,\mu\text{m}$ and $\lambda$ ranges from $0.01\,\mu\text{m}^{-1}.\text{min}^{-1}$ to $40.96\,\mu\text{m}^{-1}.\text{min}^{-1}$. Time interval between steps in simulations is $\Delta t = 0.1$min. Finite-tip-number cutoff corrections to the front velocity are not included in the theoretical prediction [8]. (C) Theoretical front speed in $(\alpha, \sigma\lambda\Delta t)$ space.

#### 2.3 Sensitivity to parameters $(\alpha, \sigma, \lambda)$

We can compute the relative importance of the parameters in setting the front speed by computing the relative sensitivities:

$$S_\alpha = \frac{\partial \ln c^*}{\partial \alpha} = \frac{4\,A(\alpha) - 4\,\ln(\sigma\lambda\Delta t)}{(2+\alpha)^2}. \tag{27}$$

with

$$A(\alpha) = \ln(2-\alpha) - \frac{3}{2}\ln(4-\alpha) + \ln 2, \tag{28}$$

$$S_\sigma = \frac{\partial \ln c^*}{\partial \ln \sigma} = \frac{4}{2+\alpha}, \tag{29}$$

$$S_\lambda = \frac{\partial \ln c^*}{\partial \ln \lambda} = \frac{2-\alpha}{2+\alpha}. \tag{30}$$

A first conclusion is that the front velocity is always more sensitive to $\sigma$ than to $\lambda$:

$$\frac{S_\sigma}{S_\lambda} = \frac{4}{2-\alpha} \geq 2. \tag{31}$$

On the other hand, the relative importance of $\alpha$ and $\sigma$ depends on the value of $\sigma\lambda\Delta t$:

$$\frac{S_\alpha}{S_\sigma} = \frac{A(\alpha) - \ln(\sigma\lambda\Delta t)}{2+\alpha}. \tag{32}$$

In practice, under the constraint of our large-time hypothesis $\Delta t\sqrt{\frac{4-\alpha}{2-\alpha}\,\lambda c^*} \ll 1$ and over the relevant set of parameters, $\alpha$ is always more important than $\sigma$ (Figure SI3). This implies that $\alpha$ is the local-dynamics parameter that most effectively tunes neuronal growth, according to the theoretical model. Strikingly, $\alpha$ is also the effective tuning parameter that distinguishes class I from class IV neurons.

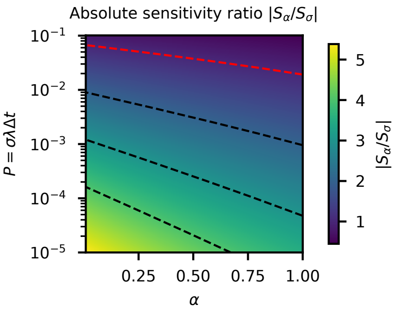


Figure SI3: Ratio $|S_\alpha/S_\sigma|$ in $(\alpha, \sigma\lambda\Delta t)$ space. The dashed red line corresponds to $|S_\alpha/S_\sigma| = 1$.

#### 2.4 Simulations of class IV neurons

Simulations of two-dimensional neurons showed that the model produces a constant-density bulk with a front propagating at constant speed (Figure SI4A and Supplementary Materials and Methods).

The one-dimensional mean-field theory predicts

$$c^*_{1\mathrm{D}} \sim \Xi(\alpha)\left(\frac{\sigma^2}{2\Delta t^\alpha}\right)^{\frac{2}{2+\alpha}} \lambda^{\frac{2-\alpha}{2+\alpha}}, \tag{33}$$

and therefore the theoretical exponents for $\sigma$ and $\lambda$:

$$B^*(\alpha) = \frac{4}{2+\alpha}, \qquad C^*(\alpha) = \frac{2-\alpha}{2+\alpha}. \tag{34}$$

The two-dimensional simulations gave front speeds lower than the one-dimensional prediction (Figure SI4B). This could be the result of several factors including the following: branch tips can disappear if they retract past a branching point, branches are not always oriented toward the front, and the finite number of leading tips can cut off the distribution of tips at the front [8].

We tested how the measured two-dimensional front speed scales with $\sigma$ and $\lambda$ at fixed $\alpha$. For each value of $\alpha$, we fitted

$$\ln c_{2\mathrm{D}} = \ln A(\alpha) + B(\alpha)\ln\sigma + C(\alpha)\ln\lambda. \tag{35}$$

The fitted exponents were larger than in the one-dimensional theory, especially at larger $\alpha$ (Table SI1, Figure SI4C).

Importantly, these two-dimensional corrections did not change the qualitative hierarchy of parameter sensitivities over the test range of parameters. In all cases, the fitted exponent of $\sigma$ was larger than that of $\lambda$, and the variation with $\alpha$ remained the dominant source of front-speed variation over the measured parameter range *in vivo*. Indeed, evaluating the fitted two-dimensional model at the mean *in vivo* values $\sigma = 0.76\,\mu\mathrm{m}$ and $\lambda = 0.029\,\mu\mathrm{m}^{-1}\mathrm{min}^{-1}$ gives an effective sensitivity to $\alpha$ $S_\alpha^{2D} \simeq 4.8$, larger than both $S_\sigma^{2D} = B(\alpha) = 1.40$–$1.69$ and $S_\lambda^{2D} = C(\alpha) = 0.445$–$0.691$. Here, $S_\alpha^{2D} \simeq \frac{\Delta \ln c_{2D}}{\Delta\alpha}$ at fixed $\sigma$ and $\lambda$, using the fitted values between $\alpha = 0.4$ and $\alpha = 1.0$.

| $\alpha$ | $n$ | $\ln A(\alpha)$ | $B(\alpha)$ | $B^*$ | $C(\alpha)$ | $C^*$ |
|---|---|---|---|---|---|---|
| 0.4 | 70 | $-5.00$ $[-5.18, -4.81]$ | 1.69 $[1.61, 1.76]$ | 1.67 | 0.691 $[0.628, 0.754]$ | 0.667 |
| 0.6 | 93 | $-4.40$ $[-4.50, -4.29]$ | 1.59 $[1.54, 1.64]$ | 1.54 | 0.615 $[0.575, 0.654]$ | 0.538 |
| 0.8 | 120 | $-3.74$ $[-3.82, -3.66]$ | 1.50 $[1.46, 1.55]$ | 1.43 | 0.521 $[0.487, 0.555]$ | 0.429 |
| 1.0 | 150 | $-3.07$ $[-3.13, -3.00]$ | 1.40 $[1.36, 1.44]$ | 1.33 | 0.445 $[0.415, 0.475]$ | 0.333 |

Table SI1: Per-$\alpha$ log-speed regressions for two-dimensional neuron simulations. The fitted model is $\ln c_{2\mathrm{D}} = \ln A(\alpha) + B(\alpha)\ln\sigma + C(\alpha)\ln\lambda$. Confidence intervals are 99% confidence intervals. The theoretical exponents are $B^* = 4/(2+\alpha)$ and $C^* = (2-\alpha)/(2+\alpha)$.

### 3 Conclusion

Together, these results provide a theoretical framework linking local branch-tip dynamics to the global expansion speed of side-branching neuronal arbors in one dimension. In contrast to classical tip-splitting branching random walks, where the front speed scales as $c^* \sim D^{1/2}r^{1/2}$, side branching leads to a distinct scaling law, $c^* \sim D^{2/3}\lambda^{1/3}$ in the diffusive case. Extending this argument to subdiffusive branch-tip motion gives

$$c^* \sim \Xi(\alpha)\left(\frac{\sigma^2}{2\Delta t^\alpha}\right)^{\frac{2}{2+\alpha}} \lambda^{\frac{2-\alpha}{2+\alpha}}, \tag{36}$$

showing that the anomalous diffusion exponent $\alpha$ directly controls the efficiency with which local tip fluctuations are converted into macroscopic arbor expansion. This formula predicts that the front speed is more sensitive to $\alpha$ than to either the fluctuation amplitude $\sigma$ or the side-branching rate $\lambda$ in one dimension.

In two-dimensional neuron simulations, the measured front speeds are lower than the one-dimensional prediction, and the fitted exponents with respect to $\sigma$ and $\lambda$ are larger than the

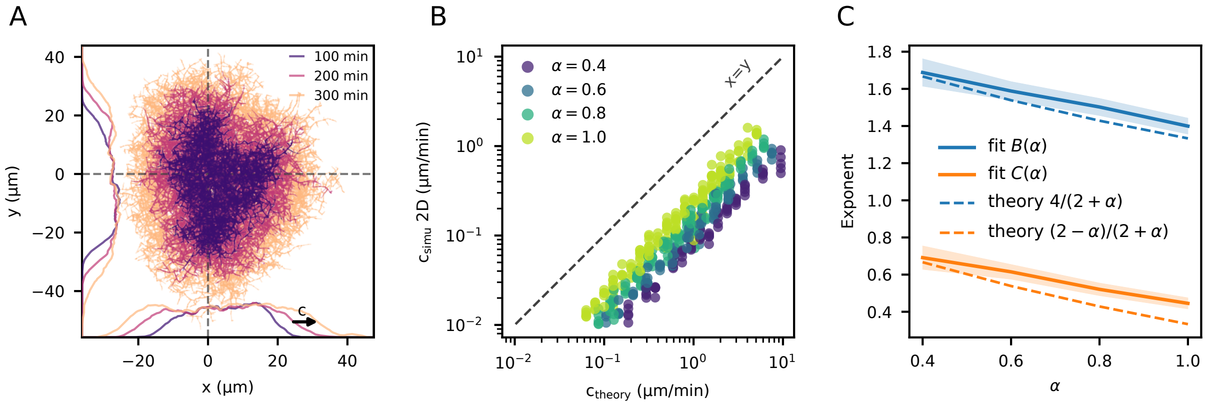


Figure SI4: (A) Overlay of ten simulations of the subdiffusive model with identical initialization. Color indicates time. Solid lines on the left and bottom show radial-density profiles along the vertical and horizontal dashed lines, respectively. The black arrow indicates front propagation. (B) Comparison of two-dimensional simulated front speeds with the one-dimensional theoretical prediction. Color indicates $\alpha$, with multiple values of $\sigma$ and $\lambda$ for each $\alpha$. Five simulations were performed per parameter combination. The dashed line indicates identity. $\sigma$ ranges from 1/4 to 4 times the mean value measured in Class IV neurons (0.76 µm), and $\lambda$ ranges from 1/4 to 32 times its mean measured value *in vivo* ($0.029\,\mu\text{m}^{-1}.\text{min}^{-1}$). (C) Fitted exponents measured in two-dimensional simulations compared with the theoretical one-dimensional exponents (blue for $\sigma$, orange for $\lambda$). The fitted exponents are larger than predicted by the one-dimensional theory, but do not change the hierarchy of parameter sensitivities over the studied range of parameters.

theoretical values in one dimension. These deviations likely reflect geometrical effects and finite-simulation corrections absent from the one-dimensional theory. Nevertheless, over the biologically relevant parameter range explored here, the hierarchy of sensitivities is preserved: the front speed remains more sensitive to $\sigma$ than to $\lambda$, and variation in the anomalous diffusion exponent $\alpha$ remains the dominant source of front-speed variation.

Thus, branch anomalous diffusion is not only the main distinguishing feature of the local growth dynamics of class I and class IV neurons, but also the most effective control parameter governing neuronal growth.

### Notations

| Symbol | Meaning | Definition / dimension |
|---|---|---|
| $\sigma$ | Standard deviation of tip increments over a lag $\Delta t$ | Measured from branch-tip displacements. |
| $\lambda$ | Lineic side-branching rate | Rate of branch creation per unit length: $[\lambda] = L^{-1}T^{-1}$. |
| $\alpha$ | Anomalous diffusion exponent | Controls the scaling of tip motion: $\alpha = 1$ is diffusive and $\alpha < 1$ is subdiffusive. |
| $D$ | Diffusion coefficient | In the diffusive limit, $D = \frac{\sigma^2}{2\Delta t}$; $[D] = L^2T^{-1}$. |
| $K_\alpha$ | Generalized diffusion coefficient | $K_\alpha \sim \frac{\sigma^2}{2\Delta t^\alpha}$; $[K_\alpha] = L^2T^{-\alpha}$. |
| $\gamma$ | Inverse decay length of the traveling front | Defined by the ansatz $u(x,t) \sim e^{-\gamma(x-ct)}$; $\gamma^*$ denotes the value that selects the front speed: $[\gamma] = L^{-1}$. |
| $c^*$ or $c^*_{1D}$ | Selected asymptotic front speed | Diffusive case: $c^* = \frac{3}{2^{2/3}}D^{2/3}\lambda^{1/3}$; subdiffusive case: Eq. 25. |
| $S_\alpha$ | Relative sensitivity of $c^*$ to $\alpha$ | $S_\alpha = \frac{\partial \ln c^*}{\partial \alpha}$. |
| $S_\sigma$ | Relative sensitivity of $c^*$ to $\sigma$ | $S_\sigma = \frac{\partial \ln c^*}{\partial \ln \sigma} = \frac{4}{2+\alpha}$. |
| $S_\lambda$ | Relative sensitivity of $c^*$ to $\lambda$ | $S_\lambda = \frac{\partial \ln c^*}{\partial \ln \lambda} = \frac{2-\alpha}{2+\alpha}$. |
| $A(\alpha)$ | Fitted prefactor in the 2D front-speed law | Defined by Eq. 35: $\ln c_{2D} = \ln A(\alpha) + B(\alpha)\ln\sigma + C(\alpha)\ln\lambda$. |
| $B(\alpha)$ | Fitted $\sigma$-exponent in 2D simulations | Coefficient of $\ln\sigma$ in Eq. 35. |
| $C(\alpha)$ | Fitted $\lambda$-exponent in 2D simulations | Coefficient of $\ln\lambda$ in Eq. 35. |
| $B^*(\alpha)$ | Theoretical $\sigma$-exponent (1D mean-field) | $B^*(\alpha) = \frac{4}{2+\alpha}$. |
| $C^*(\alpha)$ | Theoretical $\lambda$-exponent (1D mean-field) | $C^*(\alpha) = \frac{2-\alpha}{2+\alpha}$. |

Table SI2: Summary of the main parameters and derived quantities used in the supplementary analysis.
