## Supplementary material and method for "Encoding neuronal shape in the stochastic dynamics of branching processes"

May 5, 2026

[1] Aix Marseille Univ, CNRS, IBDM (UMR 7288), Turing Centre for Living Systems, Marseille, France.

[2] Aix Marseille Univ, CNRS, LAI (UMR 7333), Turing Centre for Living Systems, Marseille, France.

[3] Collège de France, 11 Place Marcelin Berthelot, Paris, France.

#### 1 Dataset

Quantifications were based on two *in vivo* time-lapse imaging datasets of dendrite morphogenesis in class I and class IV neurons.

1. **Fast image acquisition.** The first dataset consists of short movies acquired every 5 s over 10 min. It was used to quantify short-timescale tip dynamics. It includes 21 class I neurons (13 h AEL: 4 neurons, 41 tracked branches; 16 h AEL: 2 neurons, 43 tracked branches; 18 h AEL: 9 neurons, 154 tracked branches; 20 h AEL: 6 neurons, 94 tracked branches) and 16 class IV neurons (16 h AEL: 2 neurons, 29 tracked branches; 18 h AEL: 10 neurons, 433 tracked branches; 20 h AEL: 4 neurons, 318 tracked branches).

2. **Slow image acquisition.** The second dataset consists of long movies acquired every 1 min from 16 h to 21 h AEL. It was used to quantify morphodynamics at longer timescales, including global morphometrics, branching rate, tip dynamics, and contact-induced shrinkage. It includes 7 class I vpda neurons, 5 wild-type class IV neurons (4 ddaC and 1 v'ada), and 4 microtubule-depolymerized class IV neurons (all ddaC).

#### 2 Image analysis

##### 2.1 Segmentation with U-Net

Image stacks from Time lapse recordings of neurons were projected using maximum intensity projection at each time point (Figure SMM1). Dendritic arbors on the resulting images were segmented with a U-Net convolutional neural network [1] with a ResNet backbone. The network was trained to predict, for each pixel, the probability of belonging to the dendritic mask. Because dendritic pixels occupy only a small fraction of the image, training was performed with the Generalized Dice loss to compensate for class imbalance [2]. The output probability map was thresholded to obtain a binary mask of the arbor. Another U-Net was trained to segment the soma of the neuron (Figure SMM1B). The binary mask was then skeletonized to obtain a one-pixel-wide representation of the dendritic arbor. This skeleton was then converted into a graph representation (using `skan` [3] and `networkx` [4]) which was used for loop opening, branch tracking, and morphometric measurements.

##### 2.2 From a graph to a tree

**Local age regression** To estimate the local age of dendritic segments, we trained a second U-Net with a ResNet backbone to perform pixel-wise regression on the age of the arbor in the

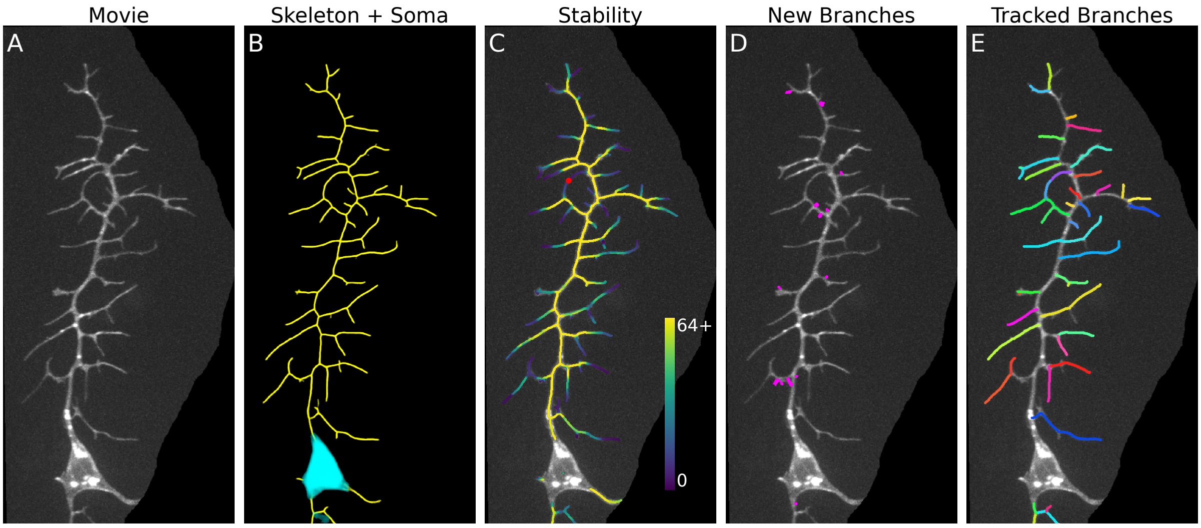


Figure SMM1: Automated image analysis. (A) Example acquisition (class I neuron). (B) Outputs of the U-Net for arbor (yellow) and soma (cyan) segmentation. (C) Stability as the mean of local age and time to disappearance (both computed with a U-Net trained on GAN-painted simulations). Color indicates stability in minutes. Each minimal loop in the skeleton is opened at the point of minimal stability (red dot) to retrieve a rooted tree structure. (D) Output of U-Net trained to detect new branches. (E) Illustration of branch tracking with custom automated tracking algorithm that leverage network structure.

last frame of a 2D+time input stack. The temporal dimension was provided as input channels, with 64 channels corresponding to 64 successive time points. Regression was performed only on pixels belonging to the segmented arbor. To correct for the imbalance in the distribution of ages, pixel-wise loss weights were assigned inversely to the frequency of each age value in the training batch. The training set consisted of GAN-painted [5] simulated neurons for which the true age of each pixel was known. In practice, the quantified local age was later combined with the time to disappearance, obtained by applying the same procedure on time-reversed movies, to define a local stability measure (Figure SMM1C).

**Loop opening** Skeletons obtained from segmented arbors could contain loops because of branch contacts and transient overlaps. Since the underlying dendritic structure is a tree, loops were removed by an iterative loop-opening procedure based on local stability. Starting from the skeleton graph, we computed a minimum cycle basis using the algorithm of [6] implemented in `networkx` [4]. For each cycle, we identified the pixel of lowest stability that was not a graph vertex (Red dot in figure SMM1C). Among all candidate pixels, the least stable one was removed, and the procedure was repeated until the graph became acyclic. This strategy used temporal information to resolve ambiguous contacts and recover a tree topology.

##### 2.3 Detection of new branches

New branches were detected with a supervised U-Net operating on pairs of consecutive frames (Figure SMM1D) [7, 8]. The task was formulated as a segmentation problem in which the network predicted the pixels belonging to nascent branches appearing between two time points. This approach was used because local deformations and branch elongation make direct frame differencing unreliable. Connected components in the predicted mask were counted as branch initiation events. The method provided the spatial position and time of appearance of newly formed branches.

#### 2.4 Tip tracking

Branch dynamics were quantified with a custom tracking algorithm, *BranchTrack*, designed for branched networks (Figure SMM1E). Tracking was formulated as a sequence of balanced linear assignment problems solved as minimum-cost matching on bipartite graphs [9, 10, 11, 12, 4]. The procedure was performed in two steps. First, each branch junction was decomposed into branch bases, defined by the junction position and the direction of each outgoing segment. Branch bases were matched between consecutive frames using a cost based on spatial distance and angular difference. Second, branch tips were matched using the tracked branch bases as anchors. For each candidate pair of tips, a cost was computed from the similarity between the paths linking each tip to its associated tracked branch base, while enforcing consistency with graph connectivity. This two-step procedure made the tracking robust to local deformations, junction sliding, and transient overlaps.

### 3 Morphometric measurements

Morphometric measurements were computed from the loop-free graph representation of each dendritic arbor. In this representation, vertices correspond to the soma, branch points, and tips, and edges correspond to dendritic segments with their ordered pixel coordinates. This tree representation was used to derive global descriptors of neuronal shape.

#### 3.1 Total length

The total length was defined as the sum of the lengths of all segments of the tree. It provides a global measure of arbor size. In practice, segment lengths were computed from the skeleton by summing Euclidean distances between successive pixels along each edge of the graph.

#### 3.2 Number of branching points

Branch points were defined as graph vertices of degree greater than or equal to three.

#### 3.3 Equivalent ellipse

To characterize the global spatial extent of the arbor, we computed an equivalent ellipse from the second moments of the tree. The dendritic arbor was treated as a planar set with uniform weight per unit length. If $N$ denotes the tree and $TL$ its total length,

$$TL = \int_N ds, \tag{1}$$

where $ds$ denotes arclength along the tree. The length-weighted centroid was defined as

$$\bar{x} = \frac{1}{TL}\int_N x\,ds, \qquad \bar{y} = \frac{1}{TL}\int_N y\,ds. \tag{2}$$

The covariance matrix of the tree was then computed as

$$\sigma_{xx} = \frac{1}{TL}\int_N (x-\bar{x})^2 ds, \qquad \sigma_{yy} = \frac{1}{TL}\int_N (y-\bar{y})^2 ds, \qquad \sigma_{xy} = \frac{1}{TL}\int_N (x-\bar{x})(y-\bar{y}) ds. \tag{3}$$

Let $\lambda_1 \geq \lambda_2$ be the eigenvalues of this matrix. The semi-major and semi-minor axes of the equivalent ellipse were defined as

$$a = 2\sqrt{\lambda_1}, \qquad b = 2\sqrt{\lambda_2}. \tag{4}$$

In the following, we used in particular the semi-minor axis $b$ as a descriptor of arbor width.

##### 3.4 Density

The area covered by the arbor was approximated by the area of the equivalent ellipse,

$$A = \pi ab. \tag{5}$$

The density was then defined as the ratio between total length and ellipse area,

$$\rho = \frac{TL}{A}. \tag{6}$$

This quantity measures how densely the dendritic length is packed within the global spatial extent of the arbor.

### 4 Statistics

##### 4.1 Correction for multiple comparisons

When multiple statistical tests were performed, $p$-values were adjusted using the Holm-Bonferroni procedure to control the family-wise error rate. This stepwise method orders the individual $p$-values from smallest to largest and compares them to progressively less stringent significance thresholds (the $k^{th}$ out of $n$ $p$-value is compared to $\frac{0.05}{n+1-k}$, all $p$-values greater than the smallest $p$-value that is above its corresponding corrected threshold are considered non-significant). The correction was applied to reduce the risk of false-positive findings that arises when several hypotheses are tested simultaneously, thereby ensuring that the overall probability of incorrectly rejecting at least one true null hypothesis remains controlled.

##### 4.2 Analysis of time-series data

Time-series data were analyzed using fixed-effects ordinary least-squares models with cluster-robust standard errors clustered at the level of the acquisition. The fixed effects control for acquisition-specific, time-invariant differences, while the cluster-robust standard errors account for the non-independence of repeated time points within the same acquisition. This approach evaluates changes across the full time course in a single model rather than testing each time point separately. Compared with separate tests at each time point, it reduces the number of statistical comparisons, limits inflation of type I error (that is, the false identification of a statistically significant effect when no true effect exists), and makes more efficient use of the temporal structure of the data. For acquisition $i$ at time point $t$, the model was

$$y_{it} = \beta_0 + \beta_1 t + \gamma_i + \varepsilon_{it}, \tag{7}$$

where $y_{it}$ is the response at time $t$ in acquisition $i$, $\beta_0$ is the reference intercept, $\beta_1$ is the common temporal slope (change per minute), $\gamma_i$ is an acquisition-specific fixed effect (constant offset), and $\varepsilon_{it}$ is the residual error. This specification estimates one shared time trend across acquisitions while absorbing baseline between-acquisition differences through fixed effects. Tests of decrease over time were based on a one-sided alternative $\beta_1 < 0$.

### 5 Quantification of branching rate $\lambda$

The lineic branching rate was quantified from the detection of new branches in the 1 min interval movies. For each frame, we divided the number of newly detected branches by the total length of the dendritic arbor at the same time point. This yielded a branching rate per unit length and per unit time for each acquisition.

For visualization only, temporal profiles were smoothed with a Savitzky-Golay filter with polynomial order 1 and window length 51 min. All statistical analyses were performed on the unsmoothed time series.
Statistics on time series were performed using a fixed-effects ordinary least-squares model (Section 4.2).
To test whether the branching rate decreased over time within each neuronal class, we fitted a cluster-robust fixed-effects ordinary least-squares model (Section 4.2). Decrease over time was assessed from the slope coefficient using a one-sided test of the alternative hypothesis that the slope is negative.
Comparisons between neuronal classes were performed on the unsmoothed time series using an cluster-robust ordinary least-squares model (Section 4.2). We tested both the difference in intercept, corresponding to a difference in branching rate level, and the difference in slope, corresponding to a difference in temporal evolution between classes.

### 6 Self-contact-induced shrinkage: quantification of $\kappa$

Self-contact-induced shrinkage was quantified from post-contact branch trajectories in the 1 min interval movies. For each annotated contact event, branch-tip displacement trajectories were aligned at the end of the contact, taken as $t = 0$. At each lag after contact, we computed the mean increment across aligned trajectories. We then defined the post-contact memory time, denoted $t_{\mathrm{mem}}$, as the first lag for which the 95% confidence interval of the mean increment contained zero for at least 3 consecutive time points. This criterion was used to identify the time at which the biased post-contact shrinkage regime ended.
We defined $\kappa$ as the mean cumulative displacement at time $t_{\mathrm{mem}}$, that is, the mean retracted length when the mean increment reached the zero-increment regime. In practice, $\kappa$ was read from the mean post-contact displacement trajectory at the class-specific value of $t_{\mathrm{mem}}$.
Confidence intervals for the mean increment and mean displacement trajectories were quantified by bootstrap resampling of post-contact trajectories with 200 bootstrap replicates. To compare $\kappa$ between classes, we used a Welch two-sample $t$-test on branch-level displacements measured at the class-specific $t_{\mathrm{mem}}$. Differences in $t_{\mathrm{mem}}$ between classes were assessed with an approximate $z$-test, using standard errors inferred from the widths of the corresponding 95% confidence intervals.

### 7 Stochastic elongation

#### 7.1 Linear-phase decomposition

Stochastic elongation was quantified from branch-tip trajectories extracted from the fast image acquisition dataset. For each tracked branch, branch length was measured over time and the resulting trajectory was decomposed into successive linear phases using piecewise linear regression following [13]. The number of breakpoints was selected by balancing goodness of fit and model complexity, using the Bayesian information criterion (Figure SMM2A). Consecutive phases with the same sign were then merged in order to match the two-state description of tip dynamics introduced in [14]. For each merged phase, the velocity was computed as the duration-weighted mean velocity of the constituent phases, and the phase duration was the sum of their durations (Figure SMM2B).

#### 7.2 Quantification of the parameters of the two-state model

Merged phases were classified as growth or shrinkage phases. For each condition, we measured the mean growth and shrinkage velocities, denoted $v_{\mathrm{on}}$ and $v_{\mathrm{off}}$, and the mean durations of

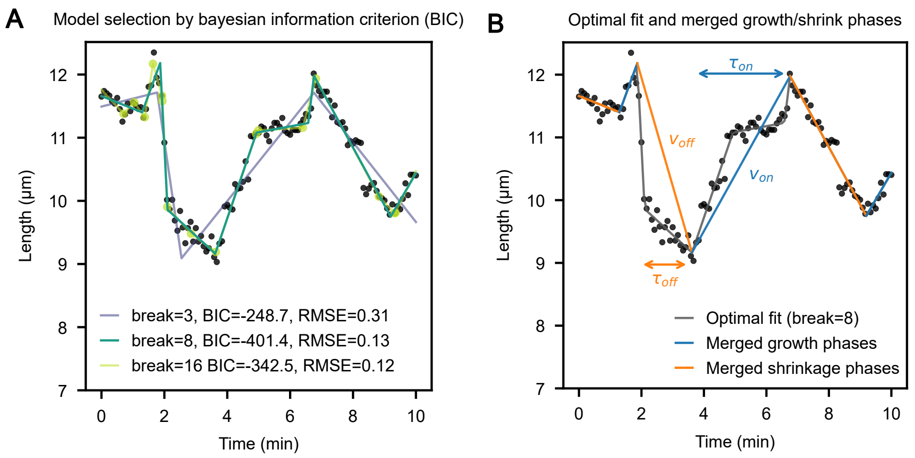


Figure SMM2: Decomposition of trajectories into phases. (A) Example of piecewise linear fit using the package in [13]. Piecewise linear functions are fitted by iteratively increasing the number of segments. The Bayesian Information Criterion (BIC) is computed to balance between error and number of break points. The fit with minimal BIC is selected to avoid overfitting. (B) Example of merged phases of the same type (growing or shrinking) from the optimal linear fit.

growth and shrinkage phases, denoted $\tau_{\mathrm{on}}$ and $\tau_{\mathrm{off}}$. The switching rates were then quantified as

$$k_{\mathrm{on}} = \frac{1}{\langle \tau_{\mathrm{off}} \rangle}, \qquad k_{\mathrm{off}} = \frac{1}{\langle \tau_{\mathrm{on}} \rangle}. \tag{8}$$

This model is also known as the asymmetric telegraph model [15]. The effective drift is

$$\bar{v} = \frac{k_{\mathrm{on}} \langle v_{\mathrm{on}} \rangle - k_{\mathrm{off}} \langle v_{\mathrm{off}} \rangle}{k_{\mathrm{on}} + k_{\mathrm{off}}}. \tag{9}$$

The effective diffusion coefficient can be computed by Green-Kubo integration of the velocity autocorrelation function of the dichotomous Markov noise derived in [16]:

$$D = \frac{k_{\mathrm{on}} k_{\mathrm{off}} (\langle v_{\mathrm{on}} \rangle + \langle v_{\mathrm{off}} \rangle)^2}{(k_{\mathrm{on}} + k_{\mathrm{off}})^3}. \tag{10}$$

##### 7.3 Dependence between phase velocity and duration (Figures 3A, 3D and S5A-H)

To quantify the dependence between phase velocity and phase duration, we fitted a linear model between the logarithm of phase duration and the absolute value of phase velocity:

$$\log \tau = \tau_0 - \frac{|v|}{v_1} + \varepsilon, \tag{11}$$

where $\tau$ is the phase duration, $v$ is the phase velocity, and $\varepsilon$ is a residual term. This regression was used to quantify the interdependence between slope and duration and to derive an alternative parameterization of stochastic elongation in which phase duration depends on phase velocity. When modeling the velocity-duration dependency, velocity was first sampled following an exponential law:

$$v \sim \mathrm{Exp}(v_{0w}), \tag{12}$$

then the logarithm of the duration according to a gaussian distribution:

$$log(\tau) \sim \mathcal{N}(\tau_{0w} - \frac{v}{v_{1w}}, \zeta_w^2), \tag{13}$$

where $w \in \{g, s\}$ indicates whether this is a growth or a shrinkage phase.

##### 7.4 Statistical analysis

Comparisons of phase velocities and phase durations between conditions were performed with Welch's $t$-tests. The resulting $p$-values were corrected within each family of tests using the Holm-Bonferroni correction (Section 4.1). To test whether the effective drift differed from zero, we used a two-sided test on the quantified drift value in each condition. For the slope-duration regression, significance of the dependence between duration and velocity was assessed from the regression coefficient associated with $|v|$.

##### 7.5 Comparison of data with independent and dependent models (Figures 3C,3E and S5G-L)

To compare the short-time behavior predicted by the two stochastic elongation models with the experimental data, we generated simulated branch-length trajectories from the independent and dependent parameterizations using the parameters quantified for each class and developmental stage. For each condition, we computed the mean displacement trajectory and its standard deviation across simulated tracks, and displayed them as a central curve with a shaded envelope. The same representation was obtained from the experimental 5 s branch tracks after aligning trajectories at their starting point. The figure therefore compares, for each condition, the envelope of real short-time branch dynamics with the envelopes predicted by the independent model, in which phase velocity and duration are sampled independently, and by the dependent model, in which phase duration depends on phase velocity. In addition, an effective diffusion coefficient was quantified from the mean squared displacement of each set of trajectories, and bootstrap resampling was used to quantify uncertainty and to test the difference between simulated and experimental envelopes.

##### 7.6 Standard deviation of 1 min increments

Tip increments were defined from the 1 min interval movies as the change in branch length between two consecutive frames. For each neuronal class and acquisition, increments from non-contacting branches were pooled within successive 50 min time bins between 16 h and 21 h AEL. Within each bin, we computed the standard deviation of the 1 min increments, denoted $\sigma$. The choice of a 1 min increment was set by the temporal resolution of the movies.
Statistical analyses for testing decrease of $\sigma$ and difference between neuronal classes were performed on acquisition-level values (Section 4.2). Under a diffusive approximation, the effective diffusion coefficient associated with the increments is

$$D = \frac{\sigma^2}{2\Delta t}, \tag{14}$$

where $\Delta t = 1$ min is the time interval between two frames.

##### 7.7 Drift of 1 min increments

We defined the drift $\mu$ as the mean 1 min increment of branch length within each 50 min time bin. For each time bin, we also tested whether $\mu$ differed from zero using one-sample $t$-tests across acquisitions. Comparisons between classes were performed with the same cluster-robust regression framework, testing both differences in level and in slope (Section 4.2).

##### 7.8 Anomalous diffusion exponent

Long-time tip dynamics were quantified from the mean squared displacement (MSD) of branch-length trajectories measured in the 1 min interval movies, which equals $\sigma^2$ in the absence of drift. For each acquisition, only non-contacting branches were retained, and trajectories were truncated to 300 min after the start of the movie. For a lag $\tau$, the MSD was computed as

$$\mathrm{MSD}(\tau) = \left\langle (X_{t+\tau} - X_t)^2 \right\rangle, \tag{15}$$

where $X_t$ denotes branch length and the average was taken over all valid time origins and trajectories.

The anomalous diffusion exponent $\alpha$ was quantified by fitting the power-law relation

$$\mathrm{MSD}(\tau) \propto \tau^{\alpha} \tag{16}$$

in log-log space, using linear regression of log(MSD) against log($\tau$). The fit was performed with lags ranging from 1 to 60 min. An exponent $\alpha = 1$ corresponds to diffusion, $\alpha < 1$ to subdiffusion, and $\alpha > 1$ to superdiffusion.

Global values of $\alpha$ were quantified for each acquisition. Comparisons between classes were performed on acquisition-level values using Mann-Whitney tests.

### 8 Simulations of class I and class IV neurons

We simulated dendritic morphogenesis in two dimensions. The neuron was represented by a tree of points in two dimensions that define a series of segments. Hereafter, and following the terminology of the SWC format [17], we refer to these points as compartments. Each compartment is defined by its own identifier, coordinates (x,y), its width, and the identifier of its parent compartment (except for the root of the tree which has no parent), such that the segment whose extremities are a compartment and its parent defines an element of a neuron branch. This representation is inspired by and allows direct export to SWC format and computation of the same morphometric descriptors as for experimental neurons (Figure SMM3). Simulations were initialized from real neurons imported from an SWC file. For each neuron in the 1 min dataset, 10 simulations were run, initialized with the shape of the neuron at 16 h AEL. The time step was set to $\Delta t = 1$ min, resulting in 300 temporal steps spanning 16 h to 21 h AEL.

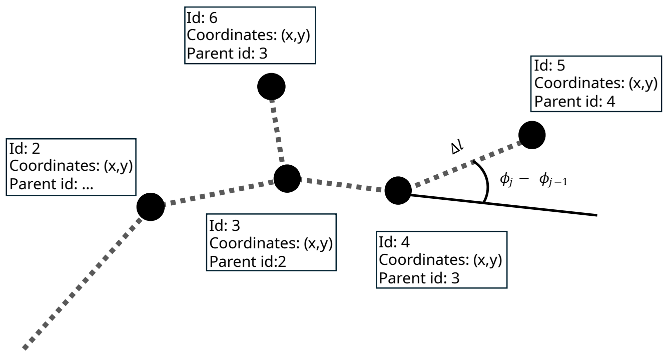


Figure SMM3: Geometric representation of neurons in simulations. The arbor is represented as a tree of compartments. Each compartment stores an identifier, its coordinates, and the identifier of its parent compartment, so that successive compartments define branch segments. When the branch grows by $\Delta l$, its angle is updated with angular diffusion following equation 17.

#### 8.1 Geometric representation of branches

Each segment was assigned a finite width and treated as a geometric object in the plane for collision detection. Branches were discretized into compartments of prescribed length and width and organized as a tree. During shrinkage, terminal compartments were removed one by one from the tip backward, and shrinkage was not allowed to proceed beyond the nearest branching point. In that case, retraction stopped at the branching point rather than deleting the parent branch.

#### 8.2 Tip growth and shrinkage models

Tip dynamics were simulated with three alternative stochastic models. In all cases, growth and shrinkage were applied at the tip of each branch.
The direction of growth was updated with angular diffusion, as in [14]. If $\phi_j$ is the angle of compartment $j$, then

$$\phi_j = \phi_{j-1} + \sqrt{\frac{2\Delta l}{l_p}}\,\eta, \tag{17}$$

where $\Delta l$ is the grown length, $l_p$ is the persistence length, and $\eta$ is a centered Gaussian random variable (Figure SMM3). This construction introduces directional persistence, with larger $l_p$ corresponding to straighter branches. $l_p$ was set to 30 µm.
During shrinkage, compartments were iteratively shrinked from the tip of the branch until the length to be retracted was consumed or a branching point was reached. The latter case meaning the disappearance of the branch tip.

In the following paragraphs we detail the three stochastic models considered for branch elongation.

##### 8.2.1 Two-states Markov model

Branches alternated between growth and shrinkage phases with switching rates $k_{\text{on}}$ and $k_{\text{off}}$, which means that durations of growth and shrinkage phases were sampled from exponential distributions: $\tau_{\text{on}} \sim \text{Exp}(k_{\text{off}})$ and $\tau_{\text{off}} \sim \text{Exp}(k_{\text{on}})$. During a growth phase, the tip advanced at constant speed $v_{\text{on}}$ until a stochastic switch to shrinkage occurred. During a shrinkage phase, the tip retracted at constant speed $v_{\text{off}}$ until a switch back to growth occurred. 1 min increments were obtained by sampling from these

##### 8.2.2 Coarse-grained diffusive model

Tip increments were generated from a diffusive process without temporal memory. At each time step, the increment was sampled independently from a Laplace distribution ($p(\Delta X = x) = 1/(2b)\exp(-|x-\mu|/b)$) with mean $\mu = 0$ (except in Figure S6) and scale $b = \sigma/\sqrt{2}$. This provided a memoryless baseline model with the correct short-time increment distribution but no temporal correlation between successive increments. $\sigma$ implicitely captures the dependency between linear phase duration and velocity (Figure 3).

##### 8.2.3 Anomalous diffusion model

To model anomalous diffusion dynamics, we considered slowly decaying correlations between tip increments (as a power law), using the ARFIMA (autoregressive fractionally integrated moving average [18]) framework. Tip increments were generated with an ARFIMA$(0, d, 0)$ process, where the fractional order of differencing $d$ is a real number that relates to the anomalous diffusion exponent $\alpha$ as follows:

$$d = \frac{\alpha - 1}{2}. \tag{18}$$

In the present case, the model contains no autoregressive term and no moving-average term, but introduces a fractional differencing parameter $d$ that generates long-range temporal correlations. Concretely, one starts from a sequence of independent and identical random variables, and convolves them with a fractional kernel whose weights decay algebraically (i.e. a power law) with the time lag: for independents $\epsilon \sim \text{Laplace}(0, 1/\sqrt{2})$:

$$\Delta X_t = \sigma \sum_{k=0}^{\infty} \frac{\Gamma(k+d)}{\Gamma(d)\,\Gamma(k+1)}\,\epsilon_{t-k}, \tag{19}$$

The result is that each increment retains a trace of all previous innovations, with more recent ones weighted more heavily but distant ones never fully forgotten. It is this slow, power-law decay of correlations that makes the process capable of generating long-range temporal correlations and, consequently, anomalous (here, subdiffusive) scaling at large times: the autocorrelation function of the ARFIMA$(0, d, 0)$ process is

$$\rho(\tau) = \prod_{j=1}^{\tau} \frac{j-1+d}{j-d}, \qquad \rho(0) = 1, \tag{20}$$

and decays as a power law with exponent,

$$\rho(\tau) \sim \frac{\Gamma(1-d)}{\Gamma(d)}\,\tau^{2d-1} \quad \tau \to \infty. \tag{21}$$

This makes ARFIMA well suited to our data because it has two experimentally measurable ingredients as explicit parameters: the 1 min increments standard deviation $\sigma$ and the anomalous diffusion exponent $\alpha$. Both play a role in short-term and long-term dynamics, but we could say that short-term dynamics is mainly modulated by $\sigma$, while long-term tip dynamics is modulated by $\alpha$.

#### 8.3 Side branching

New branches were created stochastically along existing segments. The branching rate was taken to be identical for all branches, irrespective of branch order. At each time step, each eligible compartment produced a Poisson-distributed number of branching events with mean $l_c \lambda \Delta t$, with $l_c$ the compartment length and $\Delta t$ the step of the simulation. The position of the branching event was sampled uniformly along the parent compartment. The initial length of the daughter branch was set to 0.1 µm.

The initial angle of the daughter branch was sampled relative to the local direction of the parent segment. In the implementation, daughter branches were emitted laterally, with a sign chosen uniformly on the two sides of the parent branch, and an angular deviation drawn from a von Mises distribution centered on lateral branching. If $\theta$ denotes the angle of the daughter branch relative to the parent, then

$$\theta = \pm\left(\frac{1}{2}\xi + \frac{\pi}{2}\right), \qquad \xi \sim \text{vonMises}(0, \theta_{\text{branch}}), \tag{22}$$

where the sign is sampled with probability 1/2 on each side and $\theta_{\text{branch}}$ controls the concentration of the branching angle. When $\theta_{\text{branch}} = 0$, the branching direction is broadly distributed around the lateral direction, while large values of $\theta_{\text{branch}}$ concentrate daughter branches near $\pm\pi/2$. In all simulations, $\theta_{\text{branch}}$ was set to 1.

#### 8.4 Contact-induced shrinkage

Self-contact was handled explicitly during growth. When a newly grown tip compartment intersected the existing arbor, a contact event was triggered. In the simplest implementation, contact induced a one-step backward response of the tip.

**Collision detection** Collision between a growing tip and an existing branch segment (defined by the line between a compartment and its parent) can be prohibitive in terms of computational time when done naively by testing the intersection of every tip with every segment. Collision detection was accelerated with a tile-based spatial indexing strategy. The simulation domain was partitioned into square tiles, and each compartment was assigned only to the tiles overlapped by its polygonal representation. Candidate collisions were first identified from tile co-occupancy, and exact tests were then performed only on this restricted set by polygon intersection between the growing tip compartment and existing compartments. This reduced the number of pairwise intersection tests dramatically and therefore improved computational complexity by avoiding exhaustive comparisons between all compartments at each growth step. In practice, this tile-based strategy was essential to keep simulation times tractable when running large ensembles of morphogenetic simulations from many starting arbors and replicate seeds.

#### 8.5 Temporal evolution of the parameters

The parameters of the branch program were allowed to evolve over developmental time through piecewise-constant schedules. For the diffusive and ARFIMA models, the mean and standard deviation of the increment distribution were linearly interpolated between values measured at times 25, 75, 125, 175, 225, and 275 min in the experimental movies. In the ARFIMA model, the correlation parameter $d$ was kept constant within each neuronal class, while the drift and increment scale followed these time schedules. For the two-state Markov model, the parameters $v_{\text{on}}$, $v_{\text{off}}$, $k_{\text{on}}$, and $k_{\text{off}}$ were updated at times 0, 150, and 300 min using the experimentally measured values for each class. In the two-state Markov model, zero-drift was ensured by shifting $v_{\text{on}}$ and $v_{\text{off}}$ so that the effective drift cancelled.

A time dependence of the branching rate was introduced through a piecewise schedule. In the simulations without branch-age dependence, the branching rate decreased linearly from 0.036 to 0.022 $\mu\text{m}^{-1}.\text{min}^{-1}$ between the beginning and the end of the simulation.

Contact-induced retraction parameters were also class dependent but did not depend on time in the simulations reported here. The corresponding mean retraction lengths were set to $-1.1$ µm for class I and $-1.7$ µm for class IV and class IV microtubule-inhibited neurons.

#### 8.6 Simulation outputs

Simulated arbors were exported as SWC trees with geometric and temporal annotations. Morphometric descriptors were then computed from these simulated trees, allowing comparison of total length, equivalent ellipse, arbor density, and branching structure between simulations and data. For each model and neuronal class, simulations were started from all available SWC initial conditions and repeated with 10 independent random seeds per starting arbor.

### 9 Quantification of microtubule invasion speed per branch tip

To compare microtubule expansion dynamics between class I and class IV neurons, we quantified the average rate at which the microtubules network invaded dendritic branches, normalized by the number of tips of the microtubules network. The analysis was performed on manually annotated skeleton stacks of neurons expressing fluorescently labelled tubulin. For each acquisition and each analyzed time point, the full dendritic skeleton was first converted into a graph representation. In parallel, the tubulin-rich skeleton was extracted by retaining pixels assigned to the tubulin label and skeletonizing the resulting binary mask.

For each time point $t_i$, the total length of the tubulin-rich skeleton was computed as the sum of all graph edge lengths (Figure SMM4A). The number of tubulin-rich tips, denoted $N_{\text{tip}}(t_i)$, was computed from the same tubulin-rich graph as the number of terminal graph vertices (Figure SMM4B).

For each acquisition $a$, the rate of microtubule-network expansion $v_{\mathrm{MT},a}$ was estimated by fitting a linear regression of tubulin-rich length against time.
To obtain a rate of invasion that mirrors the local dynamics of the microtubule network in each branch, we normalized this expansion rate by the mean number of tips of the microtubules network in the same acquisition,

$$\overline{N}_{\mathrm{tip},a} = \frac{1}{n_a} \sum_{i=1}^{n_a} N_{\mathrm{tip}}(t_i), \tag{23}$$

where $n_a$ is the number of analyzed time points in acquisition $a$. The average microtubule invasion speed per tip was then defined as

$$c_{\mathrm{MT},a} = \frac{v_{\mathrm{MT},a}}{\overline{N}_{\mathrm{tip},a}}, \tag{24}$$

The resulting values of $c_{\mathrm{MT},a}$ were treated as acquisition-level measurements. class I and class IV neurons were compared using a two-sample Welch's $t$-test, which does not assume equal variance between groups. For each class, we report the number of analyzed neurons, the mean and standard deviation of $c_{\mathrm{MT}}$, and the Welch test statistic and associated $p$-value (Table 15, Figure SMM4C). This analysis yielded the average microtubule invasion speed per tip used to compare microtubule expansion dynamics between class I and class IV neurons.

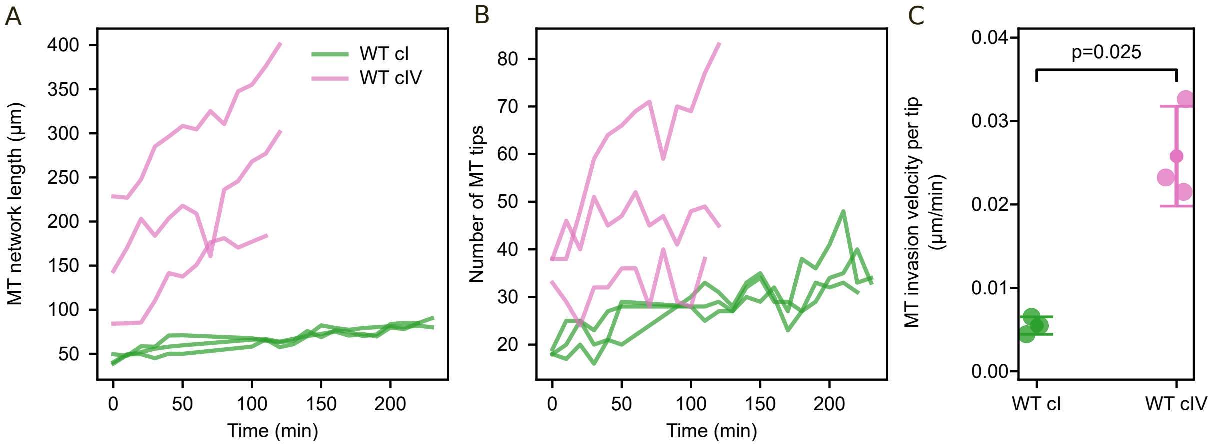


Figure SMM4: Quantification of microtubule invasion dynamics in class I and class IV neurons. (A) Total length of the tubulin-rich network in class I (green) and class IV (pink) neurons. (B) Number of tips of the tubulin-rich network in in class I (green) and class IV (pink) neurons.(C) Average microtubule invasion speed per tip, $c_{\mathrm{MT}}$, is significantly smaller in class I as in class IV neurons (4.7 fold difference, $p = 0.025$).

[7] Jake Turley, Isaac V. Chenchiah, Paul Martin, Tanniemola B. Liverpool, and Helen Weavers. Deep learning for rapid analysis of cell divisions in vivo during epithelial morphogenesis and repair. *eLife*, 12:RP87949, 2023. Reviewed preprint, version 1.

[8] Marc Karnat, Mehdi Saadaoui, Sham Tlili, Romain Karpinski, and Jean-François Rupprecht. Inferring the location and orientation of cell divisions on time-lapse image sequences. *bioRxiv*, 2024. Preprint, version 1.
